# Live-cell mapping of organelle-associated RNAs via proximity biotinylation combined with protein-RNA crosslinking

**DOI:** 10.1101/153098

**Authors:** Pornchai Kaewsapsak, David M. Shechner, William Mallard, John L. Rinn, Alice Y. Ting

**Affiliations:** Department of Chemistry, Massachusetts Institute of Technology, Cambridge, Massachusetts, USA; Department of Stem Cell and Regenerative Biology, and Molecular and Cellular Biology, Harvard University, Cambridge, Massachusetts, USA; Broad Institute of Massachusetts Institute of Technology and Harvard, Cambridge, Massachusetts, USA; Departments of Genetics, Biology, and Chemistry, Stanford University, Stanford, CA, USA

**Author notes:** Co-first authors (Pornchai Kaewsapsak, David M. Shechner).

**Keywords:** RNA localization, Subcellular Transcriptomics, Sequencing Technologies, Peroxidase, APEX2, Horseradish Peroxidase, FISH, RNA-Seq, Mammalian Cell RNA Analysis

## Abstract

The spatial organization of RNA within cells is a crucial factor in a wide range of biological functions, spanning all kingdoms of life. However, a general understanding of RNA localization has been hindered by a lack of simple, high-throughput methods for mapping the transcriptomes of subcellular compartments. Here, we develop such a method, termed APEX-RIP, which combines peroxidase-catalyzed, spatially restricted in situ protein biotinylation with RNA-protein chemical crosslinking. We demonstrate that, using a single protocol, APEX-RIP can isolate RNAs from a variety of subcellular compartments, including the mitochondrial matrix, nucleus, bulk cytosol, and endoplasmic reticulum (ER), with higher specificity and coverage than do conventional approaches. We furthermore identify candidate RNAs localized to mitochondria-ER junctions and nuclear lamina, two compartments that are recalcitrant to classical biochemical purification. Since APEX-RIP is simple, versatile, and does not require special instrumentation, we envision its broad application in a variety of biological contexts.

## Introduction

Spatial compartmentalization of RNA is central to many biological processes across all kingdoms of life, and enables diverse regulatory schemes that exploit both coding and noncoding functions of the transcriptome. For example, the localization and spatially restricted translation of mRNA plays a fundamental role in asymmetric cell division in bacteria and yeast, body-pattern formation in *Drosophila* and *Xenopus*, signaling at mammalian neuronal synapses (Jung et al. 2014), and a wide variety of other biological contexts. In another example, the localization of noncoding RNAs (ncRNAs) can play an architectural role in the assembly of subcellular structures, including short-range chromatin loops, higher-order chromatin domains, and large sub-nuclear structures like nucleoli and Barr bodies(Rinn and Guttman 2014; Engreitz, Ollikainen, and Guttman 2016). However, despite these examples, our general understanding of the breadth and biological significance of RNA subcellular localization remains inchoate.

Techniques that elucidate the subcellular localization of RNAs are therefore critical for advancing our understanding of RNA biology. Classically, such techniques rely either on imaging or biochemical approaches. Imaging methods–such as Fluorescence *In Situ* Hybridization (FISH) and artificial RNA reporter schemes–are powerful tools for elucidating the positions of a small number of target RNAs at low-to-moderate throughput(Wilk et al. 2016; K. H. Chen et al. 2015; Paige, Wu, and Jaffrey 2011; Hocine et al. 2013; Nelles et al. 2016). Alternatively, unbiased approaches for RNA discovery couple biochemical manipulations to deep sequencing. For example, the RNA partners of proteins with characteristic subcellular localization can be identified through a variety of techniques that couple protein immunoprecipitation to RNA-seq(Ule et al. 2003; Christopher Gilbert et al. 2004). Such methods have revealed many localized mRNAs, in addition to novel non-coding RNAs involved in RNA splicing(Chi et al. 2009) and RNAi(Motamedi et al. 2004). On a broader scale, a deep sampling of RNAs residing within a cellular compartment—for example, an intact organelle of interest, or partitions along a sucrose gradient, can be identified by coupling subcellular fractionation to RNA-Seq (“Fractionation-Seq”)(Sterne-Weiler et al. 2013; Mercer et al. 2011).

However, a technological gap exists among these current methods for studying RNA localization. Imaging approaches are of limited throughput, and may require specialized reagents, constructs, or microscopes that are only accessible to a handful of laboratories (Wilk et al. 2016; K. H. Chen et al. 2015; Paige, Wu, and Jaffrey 2011; Hocine et al. 2013; Nelles et al. 2016). The efficacy of immunoprecipitation-based approaches is highly sensitive to the antibodies and enrichment protocols used (Hendrickson et al. 2016) and captures only RNAs that are directly complexed with each target protein. Fractionation-Seq is applicable only to organelles and subcellular fractions that can be purified, and is frequently complicated by contaminants (false positives) and loss of material (false negatives) Therefore, a new technology is needed for unbiased and large-scale discovery and characterization of RNA *neighborhoods,* with high spatial specificity, and within cellular structures that cannot be enriched by biochemical fractionation.

Here we introduce such a technology–termed APEX-RIP– that enables unbiased discovery of endogenous RNAs in specific cellular locales. APEX-RIP merges two existing technologies: APEX (engineered ascorbate peroxidase)- catalyzed proximity biotinylation of endogenous proteins(Rhee et al. 2013), and RNA ImmunoPrecipitation (RIP)(Christopher Gilbert et al. 2004). We demonstrate that APEX-RIP is able to enrich endogenous RNAs in membrane-enclosed cellular organelles, such as the mitochondrion and nucleus, and in membrane-abutting cellular regions such as the cytosolic face of the endoplasmic reticulum. The specificity and coverage of this approach are much higher than those obtained by traditional Fractionation-Seq. Moreover, by applying APEX-RIP to multiple mammalian organelles, we have generated high quality datasets of compartmentalized RNAs that should serve as valuable resources for testing and generating novel hypotheses pertinent to RNA biology.

## Development of APEX-RIP method and application to mitochondria

APEX is an engineered peroxidase that can be targeted by genetic fusion to various subcellular regions of interest(Rhee et al. 2013) (Figure 1A). Upon addition of its substrates, biotin-phenol (BP) and hydrogen peroxide (H_2_O_2_), to live cells, APEX catalyzes the formation of biotin-phenoxyl radicals that then diffuse outward and covalently biotinylate nearby endogenous proteins. More distal proteins are not significantly labeled because the biotin-phenoxyl radical has a half-life of less than 1 millisecond(Wishart and Madhava Rao 2010). Previous work has shown that APEX-catalyzed proximity biotinylation, coupled to streptavidin enrichment and mass spectrometry, can generate proteomic maps of the mitochondrial matrix, intermembrane space, outer membrane, and nucleoid, each with <5 nm spatial specificity(Rhee et al. 2013; Hung et al. 2014; Hung et al. 2017; Han et al. 2017).

**Figure 1.**
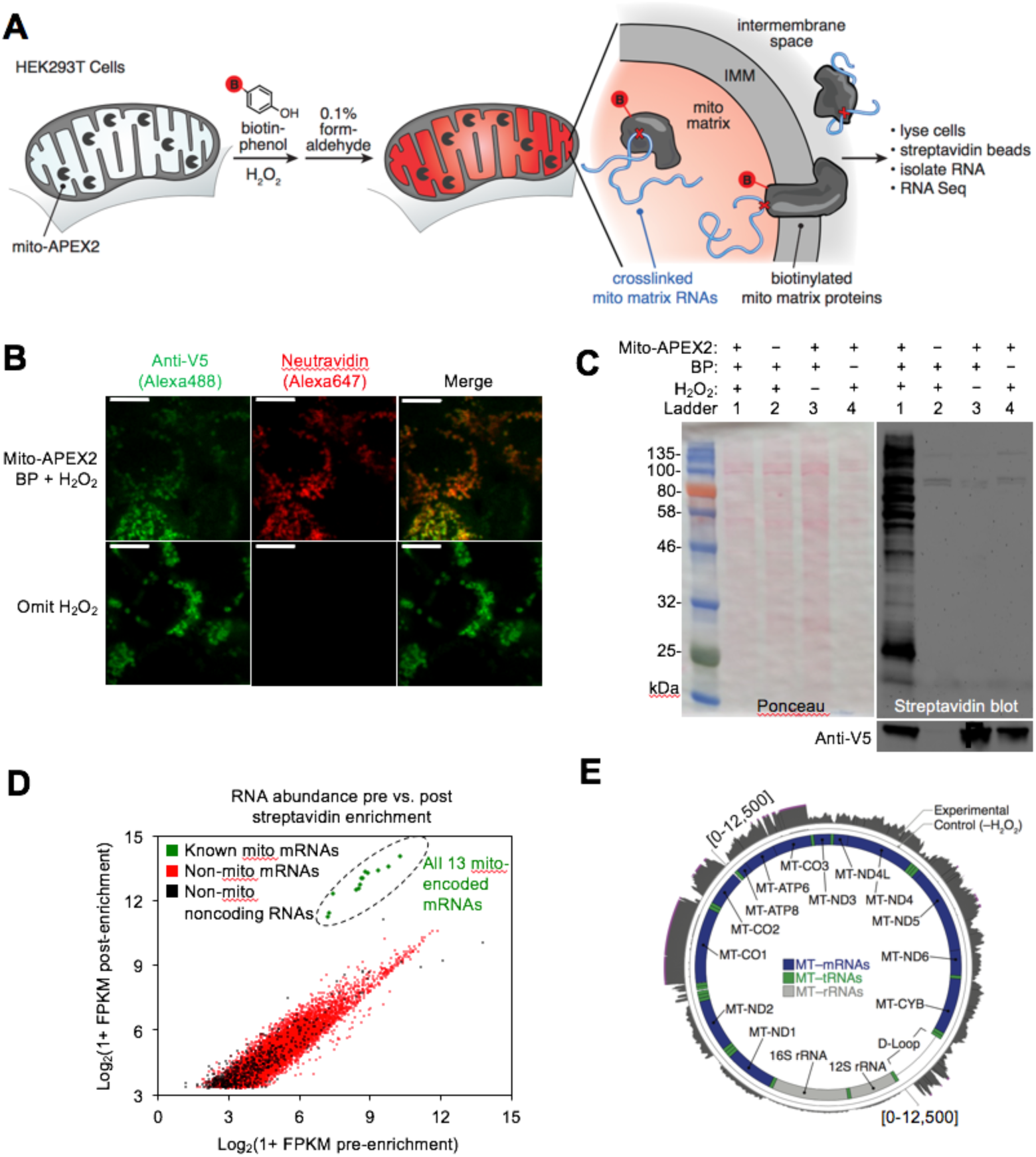
APEX-RIP in mitochondria. (**A**) Overview of the APEX-RIP workflow. Cells expressing APEX2 (grey ‘pacmen’) targeted to the compartment of interest (here, the mitochondrial matrix.) are incubated with the APEX substrate biotin-phenol (BP; red B: biotin). A one-minute pulse of H_2_O_2_ initiates biotinylation of proximal endogenous proteins(Rhee et al. 2013), which are then covalently crosslinked to nearby RNAs by 0.1% formaldehyde. Following cell lysis, biotinylated species are enriched by streptavidin pulldown, and coeluting RNAs are analyzed by RT-qPCR or RNA-Seq. IMM: inner mitochondrial membrane. (**B**) Imaging APEX2 biotinylation *in situ*. HEK 293T cells expressing V5-tagged mito-APEX2 were biotinylated and fixed as described in (A) and stained as indicated. The bottom row is a negative control in which H_2_O_2_ treatment was omitted. Scale bars, 10 μm. (**C**) Streptavidin blot analysis of whole cell lysates prepared as described in (A). *In situ* biotinylation (lane 1) is ablated in the absence of the APEX2 protein, H_2_O_2_, or BP. Anti-V5 blot detects expression of mito-APEX2. (**D**-**E**) mito-APEX-RIP efficiently recovers the mitochondrial transcriptome. (**D**) Gene-level RNA-Seq analysis of mito-APEX-RIP enrichment for RNAs longer than 200nt. Data from one biological replicate are shown. (**E**) Nucleotide-level RNA-Seq analysis of mito-APEX-RIP, mapped to the human mitochondrial genome (innermost circle). Outermost circle: reads from the full APEX-RIP protocol; middle circle: reads from the negative control. Note the enrichment of several itochondrially-encoded tRNAs and the D-loop leader transcript. Ribosomal RNAs were removed during library preparation (*see methods*). See also: Figure S1.

Because most cellular RNAs exist in close proximity to proteins, we reasoned that APEX-tagged subcellular proteomes could also provide access to the nearby RNA content, if proteins and RNA could be crosslinked together *in situ*, immediately before or after APEX labeling. As our first target organelle, we selected the mitochondrion because its RNA content--derived from both the mitochondrial genome and from imported, nuclear-encoded RNAs--has been extensively characterized by a wide array of complementary methods (Mercer et al. 2011; Alán et al. 2010; Piechota et al. 2006; Ro et al. 2013), hence providing a “gold-standard” to which we can compare our results. The mitochondrial matrix was also the first mammalian compartment mapped by APEX proteomics methodology(Rhee et al. 2013). As a RNA-protein chemical crosslinker, we opted for mild formaldehyde treatment, which covalently captures most protein-protein and protein-nucleic acid interactions, and can be achieved with minimal disruption of native interactions in live cells. It is for these reasons that formaldehyde is used for several RIP(Chris Gilbert and Svejstrup 2006) technologies for identifying the RNA partners of specific proteins of interest, including our own “fRIP-Seq” protocol (Hendrickson et al. 2016).

Since it was unclear *a priori* whether APEX-catalyzed biotinylation should precede or follow the formaldehyde crosslinking step, we explored both schemes in parallel (Figure S1A; *see methods*). Each protocol, applied to HEK 293T cells expressing mitochondrially-localized APEX2 (“mito-APEX2”, Figures 1B-C), resulted in clear enrichment of fifteen mitochondrial-encoded RNAs—relative to the cytosolic marker *GAPDH*—as gauged by RT–qPCR (average of 49.3±3.5 and 60.9±4.1-fold enrichment, respectively, Figure S1A). Assuming that fixing cells prior to biotinylation would better capture transient or weak RNA–protein interactions, we selected the crosslinking-then-BP protocol for RNA-Seq analysis. While this confirmed that mitochondrial mRNAs were enriched, a sizeable “shoulder” of conspicuous off-target RNAs were also unexpectedly enriched (Figure S1B). Thus, we re-examined our labeling and crosslinking protocols, using a sampling of these off-target RNA markers (e.g., the abundant nuclear RNA *XIST*, and cytosol-localized RNAs *HOOK2* and *MAN2C1*). This more comprehensive analysis revealed that APEX labeling followed by crosslinking provides superior specificity (Figure S1C). We hypothesize that the mild formaldehyde treatment compromises membrane integrity(Fox et al. 1985), allowing BP radicals to escape to adjoining compartments when APEX labeling is performed after formaldehyde treatment.

We used the optimized APEX followed by crosslinking protocol to map mitochondrial RNAs in mito-APEX2-expressing HEK 293T cells (Figure 1D, Table 1, tab 2). Gene-level analysis, comparing RNA counts before and after streptavidin enrichment, revealed that all 13 mRNAs encoded by the mitochondrial genome were highly enriched (greater than 3.5 fold) in three independent replicates (Figures 1D and S1E, Table 1 tab 1). Enrichment was absent in negative controls with H_2_O_2_ omitted (Figure S1F). Read density plots mapped to the mitochondrial genome demonstrated that most of our captured RNAs correspond to fully-processed transcripts, including mRNAs, interstitial tRNAs, and the D-loop leader sequence from which mitochondrial transcription initiates (Figure 1E). Intriguingly, mitomRNA read densities appeared to correlate with previous measures of mRNA halflife(Nagao, Hino-Shigi, and Suzuki 2008). For example, mRNAs encoding MTCO1-3 have longer half-lives, and more reads from APEX-RIP, than mRNAs encoding MTND1-2.

## APEX-RIP mapping of the nuclear-cytoplasmic RNA distribution

Having established that APEX-RIP is both specific and sensitive in the mitochondrion, we next turned our attention to a more challenging compartment: the mammalian nucleus. The nucleus is more complex and has a less defined transcriptome than the mitochondrial matrix, but previous Fractionation-Seq datasets, including by ENCODE(Dunham et al. 2012) (Figures S2A–D), again provide a reference list to which we can compare our results.

We generated HEK 293T cells that stably express APEX2 in the nucleus (APEX-NLS) or in the cytosol (APEX-NES; NES is a nuclear export signal). The specificity of *in situ* biotinylation by these constructs within each compartment was confirmed by imaging (Figure 2A). Whole cell lysates prepared from each cell line also produced distinct “fingerprints” of biotinylated proteins, as assayed by streptavidin blotting (Figure S1D).

**Figure 2.**
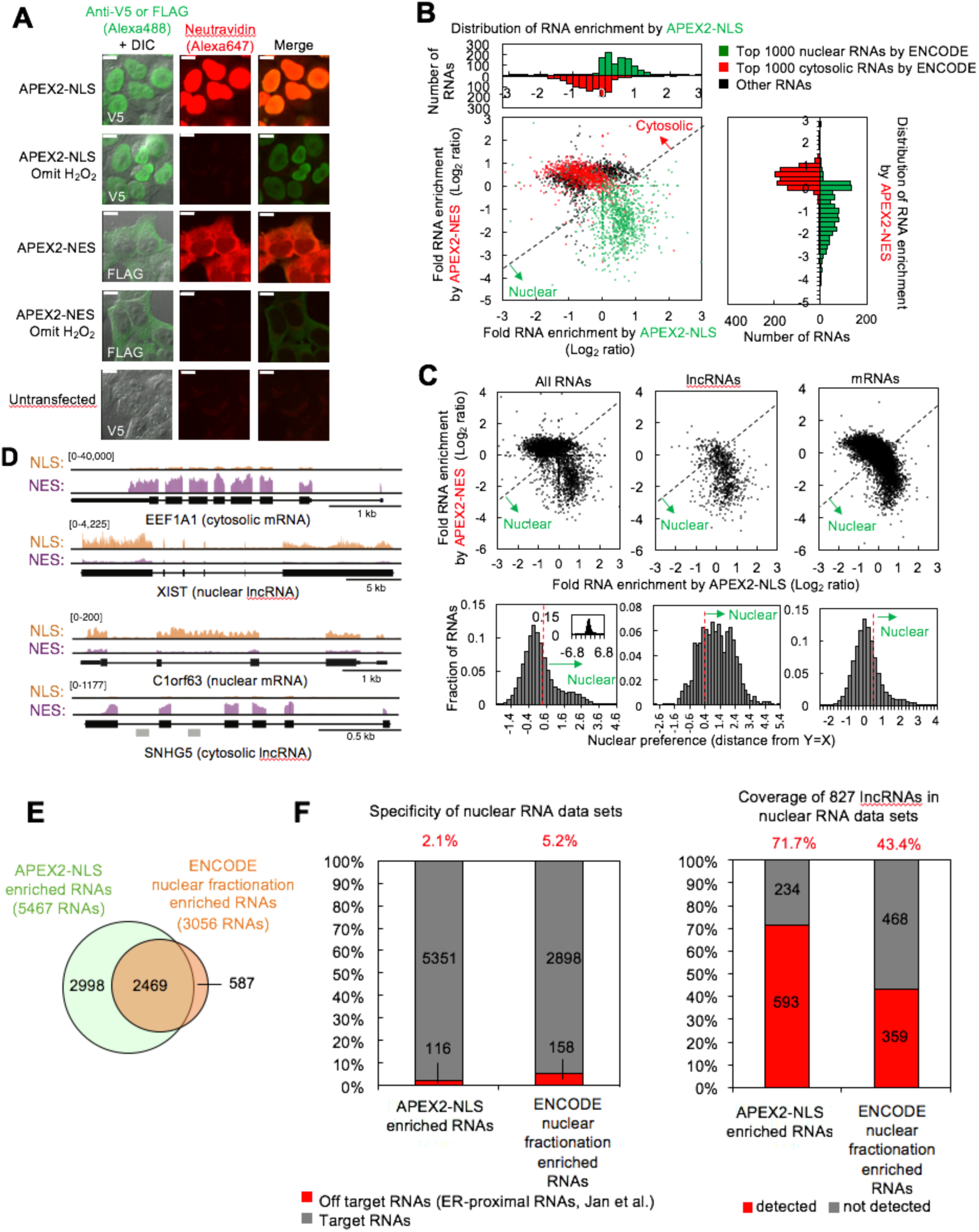
APEX-RIP mapping of the nuclear-cytoplasmic RNA distribution. (**A**) Fluorescence imaging of nuclear and cytosol-targeted APEX2 fusion constructs. HEK 293T cells expressing the indicated constructs (“NLS,” nuclear localization signal; “NES,” nuclear export signal) were labeled with biotin-phenol, crosslinked and stained as indicated. DIC, Differential Interference Contrast. Scale bars,10 μm. (**B**) Combined analysis of APEX2-NLS and NES experiments distinguishes nuclear and cytoplasmically localized RNAs. Fold enrichment values were calculated relative to matched input samples; the median values of three replicates are shown (see *methods*). The 1000 RNAs with the highest predicted nuclear and cytosolic localization by ENCODE(Dunham et al. 2012) are colored green and red, respectively (*see methods*). Histogram plots, summarizing the separation of these RNA standards by each APEX2 construct, are projected along the axes of the scatter plot and use the same scales. The black dotted line marks the cutoff between nuclear and cytosolic RNAs. (**C**) APEX-RIP captures the established nuclear-cytoplasmic distribution of mRNAs and lncRNAs. *Top*: APEX2-NLS versus APEX2-NES scatter plots, as in **(B**), for all RNAs (*left*), lncRNAs (*middle*), and mRNAs (*right*). Data are the medians of three replicates. Dotted lines mark the cutoff between nuclear and cytosolic RNAs, as in (**B**). *Bottom:* histogram plots of nuclear preference scores (*see methods*) for each class of RNA. Dotted red lines: the ROC-derived significance threshold (*see methods*). Inset: the complete distribution. (**D**) Read density plots of RNAs with stereotypical and atypical localization. For each gene, a common y-scale is used for all read tracks. SnoRNAs encoded in the *SNHG5* gene body are indicated as gray rectangles. (**E**) Venn diagram comparing APEX-RIP and ENCODE nuclear RNA datasets(Dunham et al. 2012). (**F**) Nuclear APEX-RIP is more specific and sensitive than is biochemical fractionation. *Left:* Specificity of the APEX-RIP and ENCODE nuclear RNA datasets(Dunham et al. 2012). Off-target RNAs were defined actively translated ER-proximal mRNAs (Jan, Williams, and Weissman 2014). *Right:* Recall of nuclear standard RNAs, defined as a set of 827 lncRNAs annotated by GENCODE hg19 with average pre-enrichment FPKM ≥ 1.0. See also: Figure S2.

We performed APEX-RIP on both APEX-NLS and APEX-NES cells, using the biotinylation-first/crosslinking-second protocol established above, with an additional one-minute radical-quenching step in between the APEX and crosslinking steps (Figure S3A; see *methods*). Encouragingly, “gold standard” nuclear and cytosolic RNAs (defined from the ENCODE data as the top 1000 RNAs in each compartment; see Table 2 tab 4) were enriched from the corresponding cell lines as predicted (Figure 2B *histograms* and Figures S2E–F). Moreover, when directly comparing the fold-enrichments from each compartment to one another, it was apparent that APEX-NLS had effectively enriched known nuclear-localized RNAs, while APEX-NES had enriched known cytosol-localized RNAs (Figure 2B scatter plot, Table 2 tab 3). We calculated for each RNA a “nuclear preference score,” defined as the minimum geometric distance of each point to the line y=x (corresponding to the set of genes which are not preferentially enriched from either compartment).

Receiver Operator Characteristic (ROC) analysis of these nuclear preference scores was used to filter the data and obtain final transcript lists of 5,467 nuclear RNAs and 10,130 cytosolic RNAs from living HEK 293T cells (Table 2 tabs 1 and 2). The false discovery rates of these two lists are <0.6% and <0.4%, respectively.

When plotted by nuclear preference score, the human transcriptome displayed an overall bimodal distribution, wherein the majority of species were cytoplasmic, appended by a smaller right-shifted populace of predominantly nuclear RNAs (Figure 2C, *left*). As might be predicted(Derrien et al. 2012), many of this latter group were lncRNAs, which clearly showed preferential nuclear localization (Figure 2C, *middle*). Most mRNAs appeared to be cytosolic in our data (Figure 2C, *right*). Notably, we also observed sizeable populaces of RNAs exhibiting noncanonical nuclear–cytoplasmic partitioning (Figure 2D). 3323 mRNAs–including *C1orf63*, for example (Figure 2D)–appeared preferentially nuclear. Many of these species have been proposed to play a role in dampening gene expression noise(Bahar Halpern et al. 2015). Conversely, 234 lncRNAs appeared preferentially cytoplasmic; these include the known cytoplasmic lncRNA *SNHG5*, a modulator of staufen-mediated decay that influences colorectal tumor growth(Derrien et al. 2012; Damas et al. 2016) (Figure 2D).

Our APEX-RIP nuclear and cytosolic RNA lists provide an opportunity for a head-to-head comparison with the traditional Fractionation-Seq method for mapping subcellular RNA localization. ROC analysis of the ENCODE Fractionation-Seq data yielded a list of 3,056 RNAs enriched by nuclear fractionation (Table 2 tab 5). Of these RNAs, 81% (2469) were also enriched in our APEX-RIP nuclear dataset, implying general agreement between the two technologies (Figure 2E). Notably, APEX-RIP also enriched nearly 3000 additional transcripts. These may be nuclear-localized RNAs that were opaque to the ENCODE protocol, or contaminants enriched by APEX-RIP. To address this possibility, we examined each dataset for conspicuous non-nuclear contaminants: RNAs that are known to be localized at the Endoplasmic Reticulum(Jan, Williams, and Weissman 2014). Satisfyingly, the APEX-RIP nuclear dataset, though larger, contained fewer ER contaminants than did the analogous fractionation-based dataset, implying that APEX-RIP produces higher specificity than Fractionation-Seq (Figure 2F, *left*).

To compare the coverage/sensitivity of each method (sometimes termed recall), we examined the enrichment in each dataset of lncRNAs, which are thought to be predominantly nuclear(Derrien et al. 2012). We assembled a list of 827 annotated lncRNAs (GENCODE hg19) with average FPKM pre-enrichment greater than 1.0 (Table 2 tab 7). Of these lncRNAs, 71.7 % are enriched in our APEX-RIP-derived nuclear dataset, while nuclear Fractionation-Seq from the same cell line enriched only 43.4 % (Figure 2E, *right*). We conclude that APEX-RIP can be superior to Fractionation-Seq in terms of *both* specificity and coverage, for analysis of endogenous RNA subcellular localization.

## Enrichment of RNAs proximal to the ER membrane

The above efforts establish that APEX-RIP can enrich RNAs in membrane-enclosed cellular compartments. We next sought to address whether the technique could successfully capture the transcriptomes of “open” subcellular regions. Previous proteomic work has shown that APEX tagging exhibits sufficient spatial specificity for such open compartments, since this technology has produced highly specific proteomic maps of, for example, the mammalian neuronal synaptic cleft(Loh et al. 2016), outer mitochondrial membrane(Hung et al. 2017), mitochondrial nucleoid(Han et al. 2017), and G-protein coupled receptor interaction network(Lobingier et al. 2017; Paek et al. 2017). We were unsure, however, if the additional formaldehyde crosslinking step would preserve or blur the estimated <10 nanometer spatial resolution of APEX labeling(Rhee et al. 2013).

As a test case for the generality of APEX-RIP at such open compartments, we selected the Endoplasmic Reticulum (ER). The ER is an appealing target for several reasons. First, it is host to a known set of characteristic RNAs that we can use as positive controls—the so-called “secretome”—which comprises mRNAs encoding secreted, glycosylated, and/or transmembrane proteins, that are translated by ribosomes on the rough ER. Second, the ER provides the opportunity to compare the efficacy of APEX-RIP to alternative approaches, since RNAs in this subcellular locale have been previously characterized both by Fractionation-Seq, and by a newer methodology termed proximity-dependent ribosome profiling(Jan, Williams, and Weissman 2014; Williams, Jan, and Weissman 2014). This latter technique maps active protein translation at the ER membrane by combining ribosome profiling(Ingolia et al. 2009) with proximity-restricted sequence-specific biotinylation, using an ER-targeted biotin ligase and ribosomes that are tagged with the peptide substrate (AviTag) of that ligase.

Since it was initially unclear which face of the ER membrane (cytosolic or luminal) would be most amenable to the APEX-RIP method, we generated fusion constructs that localized the peroxidase catalytic center to each (Figures 3A–B). ERM-APEX2 targets APEX2 to the ER cytosolic surface via a 27-amino acid fragment derived from the native ER membrane (ERM) protein cytochrome P450 C1. HRP-KDEL targets horseradish peroxidase (HRP) to the ER lumen via an N-terminal ER-targeting signal and a C-terminal KDEL ER-retention motif(Martell et al. 2012). We have shown that HRP catalyzes the same proximity-dependent biotinylation chemistry as APEX2(Loh et al. 2016), but has higher specific activity than APEX2 in the ER lumen(Lam et al. 2014). We generated HEK 293T cells stably expressing ERM-APEX2 and HRP-KDEL, and confirmed by microscopy and streptavidin blotting that each produced the expected labeling patterns (Figures 3C and D). Next, we compared the efficacy of each construct for target RNA isolation, using the biotinylation-first/crosslinking-second APEX-RIP protocol, and analyzing our results via RT-qPCR analysis of established secretome and non-secretome mRNAs^24^. Parallel experiments with APEX2-NES cells served as negative controls (Figure 3E).

**Figure 3.**
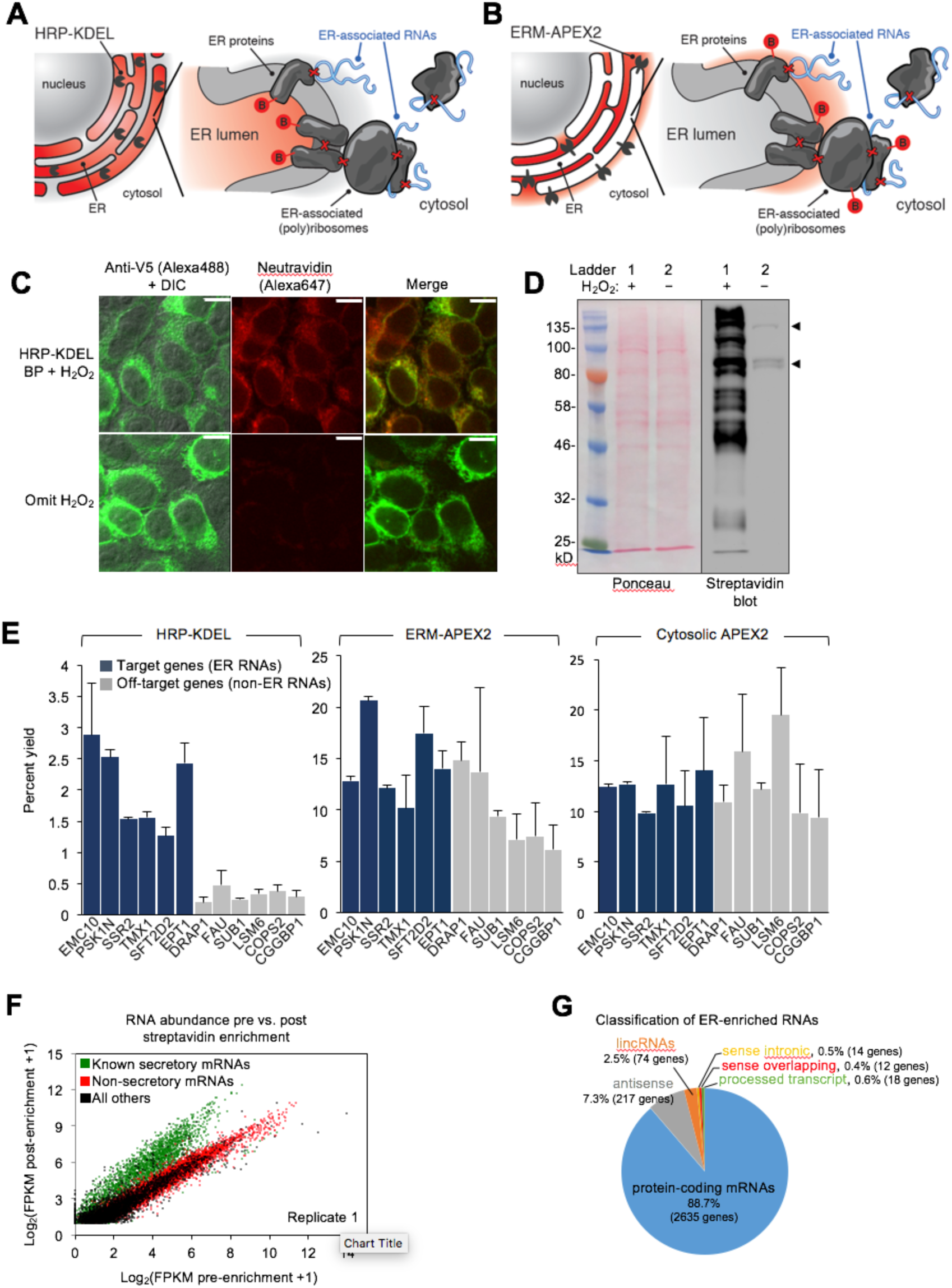
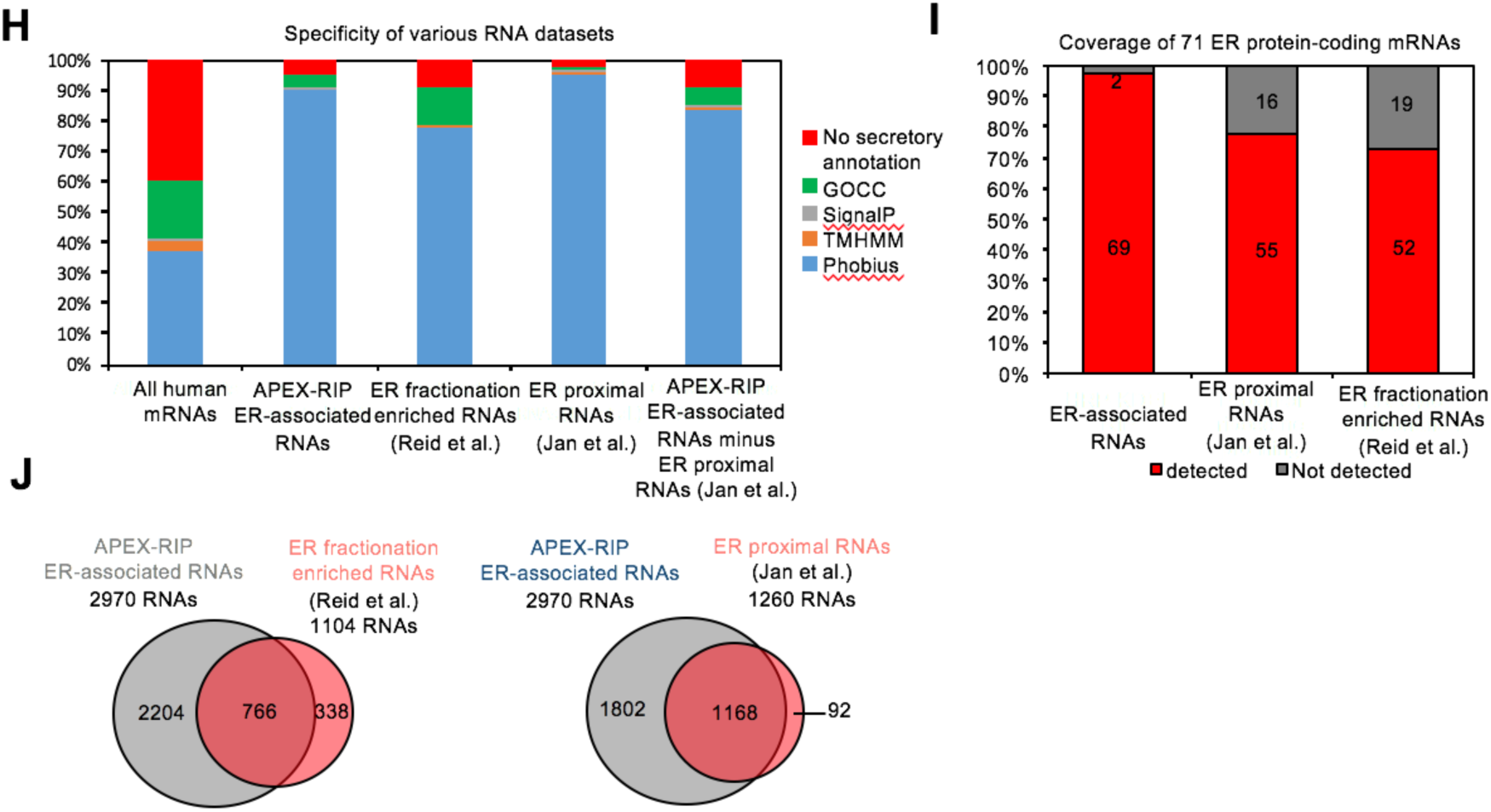
APEX-RIP maps RNAs proximal to the Endoplasmic Reticulum. (**A**—**B**) Schematics summarizing alternate ER-targeting strategies. (**A**) HRP, targeted to the ER with a KDEL sequence, biotinylates proteins within the ER lumen. Red B: biotin. Red Xs: chemical crosslinks induced by 0.1% Formaldehyde treatment. (**B**) APEX2, displayed on the ER membrane (ERM) by fusing it to the transmembrane segment of rabbit P450 C1, faces the cytosol. (**C**) Imaging HRP-KDEL-catalyzed biotinylation. HEK293T cells stably expressing HRP-KDEL were labeled with BP, fixed and imaged as in Figure 1B. DIC, Differential Interference Contrast. Scale bars, 10 μm. (**D**) Streptavidin blot detection of resident ER proteins biotinylated by HRP-KDEL, as in Figure 1C. Arrowheads denote endogenously biotinylated proteins(Chapman-Smith and Cronan 1999). (**E**) RT-qPCR analysis, comparing specificities of the labeling schemes shown in (**A–B**). Target and off-target genes were selected using previously-reported fold enrichments at the ER membrane (Jan, Williams, and Weissman 2014). Data are the mean of three replicates, ± one standard deviation. (**F**) Scatter plot showing RNA abundance before and after streptavidin enrichment. Data shown are for one replicate. Additional replicates in Figure S3B. (**G**) Classification of APEX-RIP enriched, ER-associated RNAs. Collectively, non-coding RNAs constitute 11.3% of enriched genes (335 of 2970 RNAs). (**H**) Specificity analysis for protein-coding mRNAs in our ER-associated RNA list. 95% of the 2635 APEX-RIP ER-enriched mRNAs exhibit some form of secretory annotation (as predicted by Phobius, TMHMM, SignalP, or GOCC, *see methods*), whereas only 60.3% of all human mRNAs are similarly classified (*left*). RNAs in our dataset that were not enriched by ribosome profiling (1802 RNAs) were also predominantly secretory (90.9%). (**I**) Target recall of ER APEX-RIP exceeds those of proximity-restricted ribosome profiling (Jan, Williams, and Weissman 2014) and biochemical fractionation (Reid and Nicchitta 2012). See also: Table S5. (**J**) The gene set enriched by ER APEX-RIP largely recapitulates those enriched by alternative methods. See also: Figure S3.

Intriguingly, while APEX-RIP from HRP-KDEL cells efficiently enriched target secretome mRNAs relative to non-target controls (average fold enrichment = 19.5, paired t-test p-value = 0.00009), parallel experiments in ERM-APEX2 cells exhibited only modest, qualitative enrichment of target species (average fold enrichment = 1.49, paired t-test p-value = 0.0515). Indeed, results from ERM-APEX2 cells were nearly indistinguishable from those acquired from APEX2-NES control cells (paired t-test p-value =0.830, Figure 3E, *right*). This is surprising, since proteomic experiments in HEK 293T cells expressing the identical ERM-APEX2 construct yielded highly specific enrichment of ER-localized proteins (Hung et al. 2017).

Our data strongly imply that APEX-RIP does *not* have the same spatial specificity as peroxidase-catalyzed proteomic labeling, and may be limited by perturbations induced by formaldehyde crosslinking. However, we were highly encouraged by the data obtained with the HRP-KDEL construct. We hypothesize that APEX-RIP with this construct is effective because formaldehyde crosslinking physically couples RNAs on the cytosolic face of the ER to protein complexes that are biotinylated within the ER lumen, thereby allowing target RNAs to be enriched by streptavidin (Figure 3A). Furthermore, we observed that the target specificity of this approach could be greatly improved by addition of a one-minute radical-quenching step in between the biotinylation and crosslinking steps in our protocol (Figure S3A). We surmise that this additional step prevents residual peroxidase-generated radicals from leaking into adjoining compartments when the integrity of the ER membrane is compromised during formaldehyde treatment.

Using this improved protocol, we performed APEX-RIP on HRP-KDEL cells (Table 3 tab 2). Gene-level analysis, comparing RNA counts before and after streptavidin pulldown, revealed a distinct population of substantially enriched RNAs (Figures 3F and S3B). Encouragingly, the majority (63.4%) of secretome mRNAs (defined by ER proximal RNAs(Jan, Williams, and Weissman 2014) and Phobius predicted mRNAs with exclusion of nuclear encoded mitochondrial mRNAs, see *methods*) resided in this set, while most (97.1%) mRNAs in a test set of known non-secreted genes were not enriched, thus demonstrating the ability of APEX-RIP to isolate ER-associated transcripts from the larger population of cellular RNAs (Figure 3F). Using histogram and ROC analysis, we determined the optimal log2 FKPM significance threshold cutoff for each experimental replicate (Figure S3C; *see methods*), obtaining a final list of 2970 ERM-associated RNAs that were independently enriched in multiple experiments (Table 3 tab 1). This dataset exhibited 94% specificity, based on previous secretory annotation as defined by GOCC, SignalP, TMHMM, or Phobius(Ashburner et al. 2000; Petersen et al. 2011; Krogh et al. 2001; Käll, Krogh, and Sonnhammer 2004). Figure 3H shows that we also de-enriched mRNAs lacking such signals. Coverage was likewise exceptional (97%), as gauged by the recall of 71 literature-curated well-established ER resident proteins’ mRNAs (Table 3 tab 5; Figure 3I, *see methods*).

We next compared the ERM APEX-RIP dataset to analogous results obtained by subcellular biochemical fractionation(Reid and Nicchitta 2012), and by proximity-dependent ribosome profiling(Jan, Williams, and Weissman 2014) (Table 3, tabs 3 and 4, respectively). Encouragingly, APEX-RIP captures the majority of RNAs enriched by each of these alternative techniques (70% and 93%, respectively, Figure 3J), implying broad agreement between the different methodologies. To examine this further, we quantified the specificity and coverage of each approach, as above (*see methods*). Specificity analysis demonstrated that APEX-RIP and ribosome profiling exhibited similarly high specificity (94% and 98%, respectively). However, Fractionation-Seq was substantially noisier, such that only 90% of enriched mRNAs bore a secretory annotation (Figure 3H); the remaining 10% comprised sizeable populations of conspicuous contaminants (Figure S3E). The coverage of ER-localized mRNAs retrieved by APEX-RIP (97%) was also considerably higher than those retrieved by both Fractionation-Seq and ribosome profiling (73% and 77%, respectively, Figure 3J). We attribute the enhanced coverage of APEX-RIP to its higher sensitivity, since this method appears better suited for capturing RNAs with lower abundances than do the alternative approaches (Figure S3 F-G). Such higher sensitivity may also explain why the set of RNAs enriched by APEX-RIP is so much larger than those obtained by fractionation and ribosome-profiling (Figure 3H). Excitingly, this further underscores the ability of APEX-RIP to recover RNAs that are opaque to other methods. While the vast majority (88.7%) of our enriched RNAs are mRNAs, we also enrich hundreds of noncoding RNA species–including antisense RNAs and lincRNAs (Figure 3G). These RNAs are not translated, and thus cannot be detected by ribosome profiling, and tend to be lowly expressed, making them difficult targets for either ribosome profiling or Fractionation-Seq.

In summary, APEX-RIP is superior to existing methods for mapping endogenous RNAs proximal to the ER membrane, and may be extensible to other membrane-abutting subcellular regions as well.

## Hypotheses from ER and nuclear APEX-RIP datasets

We wondered if the highly specific and comprehensive RNA subcellular localization datasets produced by APEX-RIP could be mined for new biological hypotheses. We first observed that, of the 2635 mRNAs in our ERM dataset, 141 code for mitochondrial proteins. It is thought that that the bulk of the nuclear-encoded mitochondrial proteome is translated within the bulk cytosol, or in proximity to mitochondria themselves(Lesnik, Golani-Armon, and Arava 2015), raising the possibility that the translation or subsequent processing of these 141 protein products require machinery localized to the ER. Additionally, these mRNAs may be translated at mitochondria-ER contact sites, some of which have been observed to contain ribosomes(Csordás et al. 2006). To gain initial insight into these unusual RNAs, we analyzed these 141 genes to see whether, relative to total pool of mRNAs encoding mitochondrially-localized proteins, they were enriched in particular properties (Table 4 tab 1). Intriguingly, 57.1% of these mRNAs code for transmembrane proteins (as predicted by TMHMM), compared to only 20.4% for all mitochondrial protein mRNAs (Figure 4A). Mitochondrial subcompartment analysis showed that the ER-proximal population is enriched for proteins destined for the inner mitochondrial membrane, and is depleted for resident matrix proteins, compared to the total mitochondrial proteome (Figure 4B). Interestingly, proximity-dependent ribosome profiling by Williams et al. in yeast using biotin ligase targeted to the outer mitochondrial membrane(Williams, Jan, and Weissman 2014) also showed enrichment of mRNAs encoding proteins destined for the inner mitochondrial membrane. Perhaps a subset of inner mitochondrial membrane-destined proteins are locally translated at mitochondria-ER contact sites.

**Figure 4.**
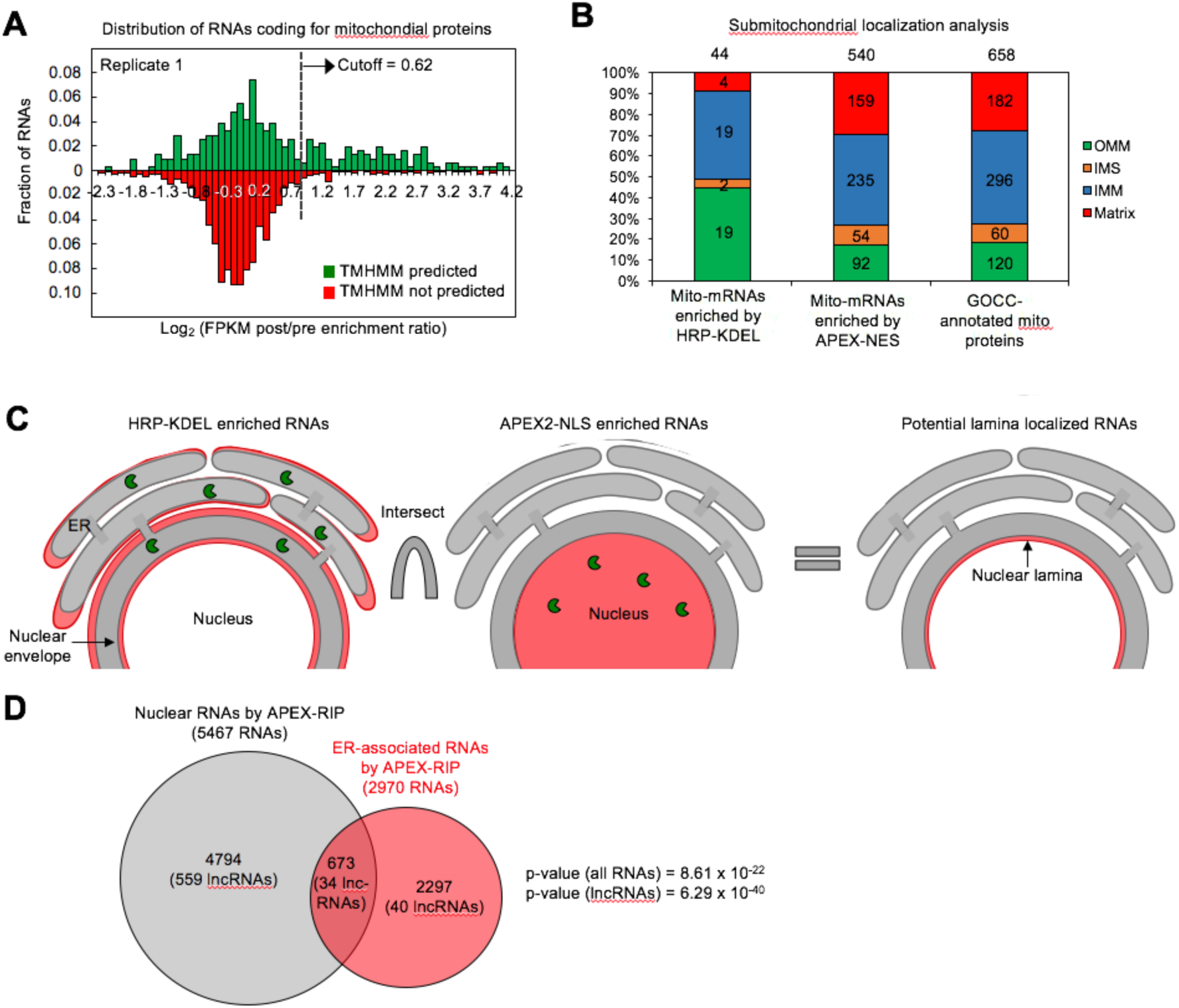
APEX-RIP reveals RNAs with potentially novel localization. (**A**) Many mitochondrial transmembrane proteins appear to be translated at the ER. mRNAs encoding mitochondrial proteins defined by GOCC and MitoCarta 1.0(Pagliarini et al. 2008; Ashburner et al. 2000), with predicted transmembrane helices (predicted by TMHMM(Krogh et al. 2001); green distribution) are preferentially enriched by HRP-KDEL APEX-RIP, relative to mitochondrial mRNAs lacking transmembrane domains (red distribution) (**B**) Predicted localization of mitochondrial proteins encoded by mRNAs that were enriched in ER- and bulk cytosol-APEX-RIP experiments (*left* and *middle*, respectively), and of all GOCC-annotated mitochondrial proteins. OMM: Outer mitochondrial membrane. IMS: Intermembrane space. IMM: Inner mitochondrial membrane. (**C**) Computational scheme for identifying putative lamina-associated RNAs. Since HRP-KDEL-enriched RNAs (*left*) comprise both ER- and lamina-associated RNAs, candidate nuclear lamina-localized RNAs, were identified as the intersection (*right*) of RNAs enriched by both APEX2-NLS (*middle*) and HRP-KDEL. Red: enriched RNAs; green pacmen: is APEX2 or HRP peroxidases. (**D**) Venn diagram identifying putative lamina-associated RNAs, defined as the overlap between HRP-KDEL- and APEX2-NLS-enriched RNAs. See also: Table 4, tab 2. The significance of overlap between ER-associated RNAs and nuclear-enriched RNAs by APEX-RIP is measured by hypergeometric test using all type of RNAs or only lncRNAs as population, respectively.

Next, we tested whether new insights could be gained by examining RNAs that APEX-RIP had enriched from more than one subcellular compartment. Because the ER lumen is contiguous with that of the nuclear envelope, we hypothesized that the HRP-KDEL APEX-RIP experiment, in addition to enriching RNAs proximal to the ER, might also enrich RNAs proximal to the nuclear membrane. This region within the nucleus, termed the nuclear lamina, is widely thought to play a critical role in gene repression(Kind and van Steensel 2010), and in shaping the global three-dimensional architecture of chromatin(C.-K. Chen et al. 2016). However, no exclusively laminar-resident RNAs have yet been identified. We hypothesized that we might identify such long-sought lamina RNAs by intersecting our APEX-RIP nuclear and ERM RNA lists (Figure 4C). Encouragingly, we observed 673 such RNAs in the intersection list, 34 of which are long noncoding RNAs (Figure 4D; Table 4 tab 2). This small list is a compelling starting point for exploration of regulatory RNAs that may reside in the nuclear lamina.

## Discussion

Methods for mapping RNA subcellular localization are constrained by the limits of their spatiotemporal precision, the diversity of RNA species that they can simultaneously analyze, their generality across cell types and compartments, and their ease of use. We believe that APEX-RIP is superior to existing imaging- and sequencing-based techniques with regard to many of these factors.

Compared to imaging-based technologies, APEX-RIP offers superior target throughput, ease of use, and temporal control. For example, although modern variants of FISH can achieve extremely high spatial precision–even enabling the visualization of individual RNA molecules(Batish, Raj, and Tyagi 2011)–this technique requires the synthesis and testing of customized fluorescent probes for each transcript under enquiry, a cumbersome process that limits throughput(Cabili et al. 2015). A highly multiplexed FISH variant, merFISH, substantially boosts throughput—enabling thousands of transcripts to be simultaneously visualized— but requires complex protocols for probe set design and imaging(K. H. Chen et al. 2015). An alternate approach, FISSEQ, achieves similar target depth without the need for gene-specific probes, but instead relies on customized instrumentation and a rococo process of *in situ* sequencing and imaging(Lee et al. 2014). Notably, without incorporating additional stains or markers, these imaging-based approaches provide little information regarding the local environment (*i.e.*, proximal cellular compartments or features) near each RNA target. Furthermore, these techniques fundamentally lack temporal precision: each requires extensively fixing and permeabilizing cells prior to data collection, during which time diffusion or the loss of cellular integrity can perturb endogenous RNA localization. This latter issue can be circumvented through a variety of live-cell imaging technologies, but these require the implementation of customized reagents that limit throughput, and may even distort the localization of the RNA targets under enquiry(Paige, Wu, and Jaffrey 2011; Hocine et al. 2013; Nelles et al. 2016). By contrast, APEX-RIP is not encumbered by any of these constraints. It does not require the development of target-specific expression constructs or probes; nor does it rely on specialized instrumentation. The ensemble of RNA targets analyzed (and, for that matter, the array of RNA classes analyzed) is theoretically limited only by the library synthesis and sequencing protocols employed. Moreover, since APEX-RIP captures only RNAs proximal to a specific subcellular compartment, and does so during a one-minute reaction, the technique offers both greater information content and higher temporal resolution than do its imaging-based alternatives.

Compared to fractionation-based technologies, APEX-RIP offers superior accuracy, ease of use, and general versatility. As illustrated in the nucleus and ER, our technique outperforms conventional Fractionation-Seq with regard to both target specificity and recall, apparently circumventing the dual issues of target loss and off-target contamination that can plague such approaches (Figures 2E-F). We ascribe this performance boost to two principal factors. First, the high spatiotemporal precision afforded by *in situ* biotinylation(Rhee et al. 2013) allows us to efficiently isolate target material from contaminants that might be difficult to remove by classical fractionation, thereby improving specificity. Second, covalently coupling target RNAs to affinity-tagged proteins allows us to recover low-abundance or weakly affiliated transcripts that might otherwise be lost during biochemical enrichment, thereby improving target recall. Perhaps more importantly, however, we have achieved these results in a variety of subcellular compartments using a common protocol, thus obviating the need to develop customized purification schemes for each compartment. This generality should enable APEX-RIP to access “unpurifiable” subcellular compartments for which such purification schemes would be impossible. While a related technology, proximity-dependent ribosome profiling, exhibits similar versatility within diverse subcellular milieus(Jan, Williams, and Weissman 2014), this approach is limited to mRNAs actively undergoing translation. It also requires biotin starvation prior to tagging, which is toxic to mammalian cells. As we have demonstrated, APEX-RIP can map diverse classes of noncoding RNA and quiescent mRNA (Figure 3G), and toxic protocols starving cells of essential nutrients for hours are not required.

The APEX-RIP methodology does have notable limitations. Cells to be analyzed must be transfected with a recombinant construct, in contrast to FISH and Fractionation-Seq, which can be performed on native tissues. APEX-RIP also gives poor spatial specificity in membrane-free subcellular regions.

The APEX peroxidase used here has also previously been used to generate contrast for electron microscopy in fixed cells(Martell et al. 2012; Lam et al. 2014), and for spatially-resolved proteomic mapping in living cells(Rhee et al. 2013; Hung et al. 2014; Loh et al. 2016; Han et al. 2017; Hung et al. 2017; Mick et al. 2015). This study extends APEX to a new class of applications and to a new biopolymer. In principle, it should be possible to utilize a single APEX-expressing cell line to characterize a target subcellular compartment by electron microscopic, proteomic, and transcriptomic means. Related methods for proteomic mapping, such as BioID(Roux et al. 2012), lack this versatility, because the underlying chemistry is not as flexible as the one-electron oxidation reaction catalyzed by APEX.

We anticipate that the initial subcellular transcriptomic map presented in this work—probing the mitochondrial matrix, cytosol, nucleus, and ER membrane of HEK293T cells—will serve as valuable resources for cell biologists. Analysis of these data has already yielded potential insight into nuclear-retained mRNAs, cytosolic lncRNAs, putative lamina-localized RNAs, and genes that may be translated locally at mitochondria-endoplasmic reticulum junctions. Applying APEX-RIP at other subcellular compartments will further expand the depth and breadth of this map. Furthermore, given the high temporal resolution of APEX-RIP, we imagine that our technology might enable profiling of subcellular RNA pools in response to acute stimuli or drugs, or throughout stages of the cell cycle and development. Collectively, such studies would yield an understanding into the biology of RNA subcellular localization at unprecedented scale.

## Significance

RNA subcellular localization is a critical factor that influences a wide array of biological processes, ranging from *Drosophila* embryogenesis to mammalian neuronal signaling. However, while this spatial layer of transcriptome regulation has been characterized in a handful of contexts, a broader understanding of its overall extent, the factors governing its establishment, and its impact on biological function, remain inchoate. The limitations hindering this understanding have been largely technical, since conventional methods—such as fluorescence *in situ* hybridization (FISH) and Fractionation-Sequencing (“Frac-Seq”)—depend upon specialized reagents and protocols that can limit throughput and general applicability. To address this fundamental need, we have developed a new strategy—APEX-RIP— which uses a simple toolkit and workflow to map the transcriptomes of discrete subcellular compartments at high depth and spatiotemporal resolution. APEX-RIP utilizes the engineered ascorbate peroxidase APEX to biotinylate proteins within a target subcellular compartment in live cells; these affinity-tagged proteins are then chemically crosslinked *in situ* to nearby RNAs. When applied to a variety of membrane-enclosed and membrane-adjacent compartments, the APEX-RIP strategy exhibited higher target specificity and coverage than do conventional fractionation-sequencing-based approaches, at a depth far exceeding those attainable by imaging-based methods. Furthermore, APEX-RIP can be applied to compartments that are recalcitrant to conventional biochemical purification. Given the superior precision, flexibility, and ease of this approach, we anticipate that APEX-RIP will provide a powerful tool for dissecting RNA subcellular localization in a broad range of biological contexts.

## Author contributions

Conceptualization, PK, DMS, AYT; Methodology, PK, DMS, AYT; Validation, PK and DMS; Formal Analysis, PK and WM; Investigation, PK and DMS; Data Curation, PK; Writing – Original Draft, PK, DMS, and AYT; Writing – Review & Editing, PK, DMS, JLR, and AYT; Visualization, PK, DMS, and AYT; Supervision, JLR and AYT; Project Administration, AYT; Funding Acquisition, JLR and AYT.

## Acknowledgements

We thank members of the Ting laboratory, especially Jeffrey Martell for valuable experimental advice and Ozan Aygun for the curated ER protein list. We thank the Rinn laboratory, especially Chinmay Shukla, for valuable computational advice. Funding was provided by the NIH (R01-CA186568 to A.Y.T. and U01 DA040612 to J.L.R.) and Stanford (to A.Y.T).

## Experimental Procedures

### Plasmids and cloning

The pCDNA3 mito-APEX plasmid was published previously (Rhee et al. 2013). The Mito-APEX2 construct was cloned from this plasmid using a two-step protocol. First, the A134P mutation(Lam et al. 2014) was introduced into the APEX gene itself, using QuikChange mutagenesis (Agilent), and thereafter the APEX2 gene was moved to the lentiviral vector pLX304 via Gateway cloning (Thermofisher), to generate plasmid pLX304 mito-APEX2.Other APEX-fusion constructs (pLX304 APEX2-NLS, pLX304 APEX2-NES, and plx304 ERM-APEX2) were cloned by Gibson assembly (NEB), using PCR to add targeting sequences and Gibson Assembly homology arms to the APEX2 gene, and joining the resulting insert into the pLX304 vector digested by BstBI and NheI. For HRP-KDEL, HRP C previously published(Martell et al. 2016) was used as a template to make HRP-KDEL-IRES-Puromycin PCR fragment. Then the insert was cloned into PCDNA3 vector digested by NotI and XbaI. Targeting sequences and restriction sites for all constructs are listed in (Table S1).

### Mammalian cell culture

Human embryonic kidney (HEK) 293T cells were obtained from ATCC, and cultured in growth media consisting of 1:1 DMEM:MEM (Cellgro), supplemented with 10% Fetal Bovine Serum (FBS), 50 units/mL penicillin, and 50 μg/mL streptomycin, at 37°C and under 5%CO_2_. Cells were discarded at 25 passages, and were periodically tested for Mycoplasma contamination using Universal Mycoplasma Detection kit (ATCC). For fluorescence microscopy imaging experiments (Figures 1B, 2A and 3C), cells were grown on 7×7-mm glass coverslips in 48-well plates. To improve cell adherence, coverslips were pretreated with 50 μg/mL fibronectin (Millipore) for 20 min at 37 °C and washed once with Dulbecco’s phosphate-buffered saline (DPBS), pH 7.4. Cells used for generating lentivirus were grown on T25 plates, in MEM supplemented as above, at 37 °C under 5% CO_2_.

### Preparation of cell lines stably expressing APEX-fusion constructs

To prepare lentivirus, one ~70% confluent T25 plate of HEK 293T cells, grown as above, was co-transfected with 2.5 μg of APEX2 fusion plasmid, along with 0.25 μg and 2.25 μg, respectively, of the lentivirus packaging plasmids VSV-G, and dR8.91(Pagliarini et al. 2008). Transfection mixes used 10 μL Lipofectamine 2000 (Invitrogen) and were brought to a final volume of 2 mL with unsupplemented MEM. The cells were transfected for 3 hours, after which media was replaced with 2 ml of fresh growth media with FBS. After 48 hours, the lentiviral supernatant was collected by aspiration and filtered through a 0.45 μm syringe-mounted filter. This filtered supernatant was immediately used to infect cells. HEK293T cells, grown in 6-well plates as described above, were infected at ~50% confluency, grown for 2 days, followed by selection in growth medium supplemented with 8 μg/mL blasticidin for 7 days, before further analysis.

For the cells stably expressing HRP-KDEL, HEK293T cells at ~60% confluency, grown in 6-well plates as described above, were transfected with the mixture of 150 μg of plasmid and 10 μL Lipofectamine 2000 (Invitrogen) in unsupplemented MEM for 3 hours, after which media was prelaced with 2 ml of fresh growth media with FBS. After 48 hours, the cells were trypsinized and replated in T25 flask in growth medium supplemented with 1 μg/mL puromycin for 7 days, before further analysis.

### *In situ* biotinylation and crosslinking

Stable-expression HEK 293T cells were grown to 90% confluency in 6-well plates, as described above. For the crosslinking–then–BP biotinylation protocol (Figure S1A, *top*), cells were washed once with 5 mL PBS, and crosslinked in 5 mL 0.1% (v/v) formaldehyde in PBS for 10 min at room temperature, with gentle agitation. The crosslinking reaction was quenched by addition of glycine (1.2 M, in PBS) to final concentration 125 mM, and gentle agitation for 5 minutes at room temperature. Crosslinked cells were then washed three times with PBS and incubated with 500 μM biotin-phenol (BP)(Rhee et al. 2013) in PBS at room temperature, for 30 min. Thereafter, H_2_O_2_ was added to a final concentration 1 mM, for 1 min. The liquid phase was then removed by aspiration, and cells were washed twice with 2 mL quenching solution (5 mM Trolox, 10 mM Ascorbate, 10 mM sodium azide, in PBS). Crosslinked, labeled cells were collected by scraping, and pelleted by centrifugation, and either processed immediately or flash frozen in liquid nitrogen and stored at – 80 °C before further analysis.

For the BP–then–crosslinking protocol (Figure S1A, *bottom*) used for mito-APEX2 experiments (Figure 1), cell growth media was replaced with fresh media supplemented with 500 μM BP. Cells were incubated in BP-supplemented media for 30 minutes at 37 °C, after which H_2_O_2_ was added to a final concentration of 1 mM. After 1 min, the media was replaced with 5 mL crosslink-quench solution (0.1% (v/v) formaldehyde, 10 mM ascorbate, and 5 mM Trolox, in PBS) for one minute, to simultaneously quench the APEX2 BP labeling reaction and initiate formaldehyde crosslinking. Thereafter, cells were washed and incubated in 5 mL of fresh crosslink-quench for two additional 1-minute incubation steps, followed by a third, 8-minute wash. Thereafter, crosslinking was terminated by the addition of Glycine, and cells were harvested as described above.

The BP–quench–then–crosslinking protocol (Figure S3A) used for all other subcellular compartments was identical to the BP–then–crosslinking protocol, except that, following BP-labling, and prior to the addition of crosslink-quench solution, cells were incubated in 2 mL azide-free quenching solution (10 mM ascorbate and 5 mM Trolox, in PBS) for one minute. Subsequently, cells were subjected to only two (1 and 9 minute) treatments in crosslink-quench solution. Thereafter, crosslinking was terminated by the addition of Glycine, and cells were harvested as described above.

### Immunofluorescence staining and microscopy

For immunofluorescence experiments (Figures 1B, 2A, and 3C), stable APEX- or HRP-expressing cells were BP-labeled and crosslinked, as above, and subsequently fixed with 4% (v/v) paraformaldehyde in PBS at room temperature for 10 min. Cells were then washed with PBS three times and permeabilized with cold methanol at – 20 °C for 5 min. Cells were washed again three times with room-temperature PBS and then incubated with primary antibodies in PBS–supplemented with 1% (w/v) Bovine Serum Albumin (BSA)–for 1 h at room temperature. After washing three times with PBS, cells were incubated with secondary antibodies and neutravidin-AlexaFluor647 (1:1000 dilution) in BSA-supplemented PBS for 30 min. Cells were then washed three times with PBS and imaged by confocal fluorescence microscopy, or in PBS at 4 °C in light-tight containers prior to imaging. Primary and secondary antibodies used were listed in Table S2.

Fluorescence confocal microscopy was performed with a Zeiss AxioObserver microscope with 63× oil-immersion objectives, outfitted with a Yokogawa spinning disk confocal head, a Cascade II:512 camera, a Quad-band notch dichroic mirror (405/488/568/647), 405 (diode), 491 (DPSS), 561 (DPSS) and 640 nm (diode) lasers (all 50 mW). Alexa Fluor488 (491 laser excitation, 528/38 emission), Alexa Fluor 568 (561 laser excitation, 617/73 emission), and AlexaFluor647 (640 laser excitation, 700/75 emission) and differential interference contrast (DIC) images were acquired through a 63x oil-immersion lens. Acquisition times ranged from 100 to 1,000 ms. For imaging quantitation and analysis, we used the SlideBook 6.0 software (Intelligent Imaging Innovations) to process and normalize the images.

The data in these figures (Figure 1B, 2A, and 3C) are representative of three independent experiments with ≥ 5 fields of view each.

### Western and Streptavidin blotting

For blotting experiments (Figures 1C, 3D and S1D), stable APEX- or HRP-expressing cells were grown in 6-well plates. After labeling, the cells were harvested by scraped, pelleted by centrifugation at 3,000×*g* for 10 min, and stored at –80 °C prior to use. Thawed pellets were lysed by gentle pipetting in RIPA lysis buffer (50 mM Tris,150 mM NaCl, 0.1% SDS, 0.5% sodium deoxycholate, 1% Triton X-100, 5 mM EDTA), supplemented with 1× protease cocktail (Sigma Aldrich),1 mM PMSF (phenylmethylsulfonyl fluoride), for 5 min at 4°C. Lysates were then clarified by centrifugation at 15,000×*g* for 10 min at 4 °C before separation on homemade 8% SDS-PAGE gels. Gels were transferred to nitrocellulose membranes, stained by Ponceau S (0.1% (w/v) Ponceau S, 5% (v/v) acetic acid, in water) for 10 min at room temperature, and imaged. The blots were then blocked with blocking buffer (3% (w/v) BSA, 0.1% (v/v) Tween-20 in Tris-buffered saline) for 1 h at room temperature, and incubated with primary antibodies in blocking buffer for 1 h more. The dilutions of the antibodies are as followed: Mouse anti-V5 antibody (Life Technologies) 1:1000 dilution and Mouse anti-FLAG antibody (Life Technologies) 1:800 dilution. Blots were rinsed four times for 5 min with wash buffer (0.1% Tween-20 in Tris-buffered saline), and then immersed in blocking buffer supplemented with Goat anti-Mouse IgG H + L-HRP Conjugate (1:3,000 dilution, Bio-Rad), for 1 h at room temperature. Blots were rinsed four times for 5 min with wash buffer, and developed with the Clarity reagent (Bio-Rad) and imaged on an Alpha Innotech gel imaging system. Processing of streptavidin blots was similar. Following Ponceau imaging, blots were blocked in blocking buffer for 30 min at room temperature, immersed in blocking buffer supplemented with streptavidin-HRP (1:3,000 dilution, ThermoFisher Scientific) at room temperature for 15 min, rinsed with blocking buffer five times for 5 min each, developed and imaged using the Clarity reagent and an Alpha Innotech gel imaging system.

The data in these experiments (Figures 1C, 3D and S1D) were also reproduced for quality control prior to quantitative PCR and sequencing.

### Streptavidin bead enrichment of biotinylated material and RNA isolation

Unless otherwise noted, all buffers used during RNA isolation were supplemented to 0.1 U/ μL RNaseOUT (Thermo Fisher), 1×EDTA-free proteinase inhibitor cocktail (Thermo Fisher) and 0.5 mM DTT, final. APEX- or HRP-expressing stable cells were grown, labeled, crosslinked and harvested as described above. Labeled cell pellets were lysed by incubation in 1 mL ice-cold RIPA buffer, supplemented with 10 mM ascorbate and 5 mM Trolox, for 5 min at 4 °C with end-over-end agitation. Samples were then sheared as described previously(Hendrickson et al. 2016) using a Branson Digital Sonifier 250 (Emerson Industrial Automation) at 10% amplitude for three 30-s intervals (0.7 s on + 1.3 s off), with 30-s resting steps between intervals. Samples were held in ice-cold metal thermal blocks throughout sonication. Lysates were then clarified by centrifugation at 15,000×*g* for 5 min at 4 °C, moved to fresh tubes and diluted with 1 mL Native lysis buffer (NLB: 25mM Tris pH 7.4, 150 mM KCl, 0.5% NP-40, 5 mM EDTA), supplemented with ascorbate and trolox), each. For each sample, 20% was removed as “input;” to the remainder was added 50 μL of streptavidin-coated magnetic bead slurry (Pierce) that had been equilibrated by two washes in 1:1 RIPA:NLB. Samples were incubated for 2 h at 4 °C with end-over-end agitation. Beads were subsequently washed with the following series of buffers (1 mL each, 5 min per wash, 4 °C, with gentle end-over-end agitation): (1) RIPA buffer, supplemented with trolox and ascorbate, (2) RIPA buffer without radical quenchers, (3) high salt buffer (1 M KCl, 50 mM Tris, pH 8.0, 5 mM EDTA), (4) urea buffer (2 M Urea, 50 mM Tris, pH 8.0, 5 mM EDTA), (5) RIPA Buffer, (6) 1:1 RIPA: NLB, (7) NLB, and (8) TE (10 mM Tris, pH 7.4, 1 mM EDTA).

Enriched RNAs were released from the beads by proteolysis in 100 μL of Elution Buffer (2% N-lauryl sarcoside, 10mM EDTA, 5mM DTT, in 1X PBS, supplemented with 200 μg proteinase K (Ambion) and 4 U RNaseOUT) at 42 °C for 1h, followed by 55 °C for 1h, as previously described(Hendrickson et al. 2016). Eluted samples were cleaned up using Agencourt RNAClean XP magnetic beads (Beckman Coulter), following the manufacturer’s 1.5 mL tube format protocol, and eluted into 85 μL H2O. Thereafter, contaminating DNA was removed by digestion with 5 U RQ1 RNase-free DNase I (Promega) in 100 μL of the manufacturer’s supplied buffer (1X final concentration) at 37 °C for 30 min. Purified RNAs were again cleaned up using Agencourt RNAClean XP magnetic beads, as above, and eluted into 30 μL H2O. The concentration and integrity of all samples was measured using an Agilent 2100 Bioanalyzer, following the “RNA Nano” or “RNA Pico” protocols, where appropriate. Samples were not heat-cooled prior to loading Bioanalyzer chips.

### Quantitative RT–PCR

For quantitative RT–PCR (qRT–PCR, Figures S1A, S1C, 3E, and S3A) RNA samples were reverse transcribed using the SuperScript III Reverse Transcriptase kit (ThermoFisher Scientific), priming with random hexamers (ThermoFisher Scientific) according to the manufacturer’s protocol. Samples were diluted with water, mixed with gene specific primers (Table S3), and Rox-normalized FastStart Universal SYBR Green Master Mix (Roche), and aliquotted into 384-well plates. qRT–PCR was performed on an Applied Biosystems 7900HT Fast real time PCR instrument, in quadruplicate. All threshold cycles (Ct, calculated per well) and efficiencies (***ε***, calculated per primer pair), were calculated from “clipped” data, using Real time qPCR Miner(Zhao and Fernald 2005). Raw C_t_ values were corrected to account for the differences in sample volume, and percent yields were calculated via the ΔCt method:

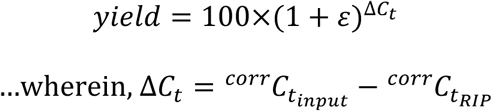

Experimental uncertainties were calculated as described previously(Shechner et al. 2015). Given D = A–B, uncertainly was calculated using the formula:

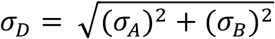

…wherein σ_A_ and σ_B_ are the measurement errors of A and B, respectively. For P, the product or quotient of values A and B, uncertainty was calculated using the formula:

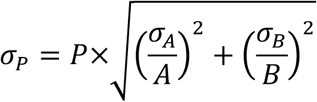

The uncertainties of other functions, *f(x)*, were calculated using the first derivative approximation:

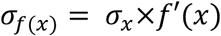

Sample sizes were determined in accordance with standard practices used in similar experiments in the literature; no sample-size estimates were performed to ensure adequate power to detect a prespecified effect size. Experiments were neither randomized nor blinded to experimental conditions. Each samples contained four replicates and no samples were excluded from analysis.

The experiments for Figures S1A, S1C, 3E, and S3A were performed once. For statistical analysis on Figure 3E, percent yield of 6 target genes were compared against percent yield of 6 non-target genes using paired t-test for both HRP-KDEL and ERM-APEX2. For comparison between ERM-APEX2 and APEX2-NES, 12 target and non-target genes were compared against each other using paired t-test.

### Library preparation, sequencing, and quantification

Purified RNA samples were depleted of ribosomal RNA using the Ribo-Zero Gold rRNA removal kit (Illumina), using 1 μL or 2 μL of Ribo-Zero rRNA Removal Solution for samples with ≤16 ng or ≥50 ng total input RNA mass, respectively, in 20 μL final reaction volume. Eluted RNA was cleaned up with Agencourt RNAClean XP beads and eluted with 19.5 μL of Elute, Prime, Fragment mix from the TruSeq RNA sample preparation kit, v2 (Illumina). Thereafter, libraries were prepared using the TruSeq RNA sample preparation kit, according to the manufacturer’s instructions, starting from “Incubate RFP” step. Each library was given a unique index during synthesis. Library concentration and quality were confirmed on an Agilent 2100 Bioanalyzer, using “DNA High Sensitivity” kits.

Indexed libraries were pooled in equimolar concentrations, with no more than ten libraries per pool, and subjected to 50 cycles of paired end sequencing, followed indexing, on two lanes of Illumina HiSeq 2500 flow cells, run in rapid mode (Genomics Core, Broad Institute of Harvard and MIT).

In general, the experiments for each construct were performed in three biological replicates. The mito-APEX experiment in Figure S1B and the mito-APEX2 negative control experiment (omit H_2_O_2_) in Figure S1F was performed in two biological replicates.

### Quantification of RNA Abundances and Folds Enrichment; Assembly of True positive and False positive lists

Deep sequencing reads were mapped to human genome assembly hg19 and UCSC known genes using TopHat2, set to default options(Kim et al. 2013). Transcript- and gene-level abundances were quantified and Cuffdiff2, set to default options(Trapnell et al. 2013), and 0.01 was added to all quantified values for ENCODE data. Enrichment analysis (*e.g.* Figures 1D, 3F, S1B, E, F, S2E, F, and S3B), was restricted to RNAs with FPKMs ≥1.0 following streptavidin pulldown. Fold enrichments were calculated as follows:

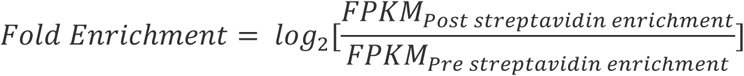

To call significantly enriched genes from our data, ENCODE data(Dunham et al. 2012), and ER-fractionation data(Reid and Nicchitta 2012), threshold cutoffs were determined using Receiver Operator Characteristic (ROC) analysis(Fawcett 2006), employing sets of true-positive and false-positive genes identified as described below. At each fold enrichment value of the data, true positive rate (TPR – fraction of detected true positive genes above the fold enrichment value) and false positive rate (FPR – fraction of detected false positive genes above the fold enrichment value) are calculated. The fold enrichment value that maximizes TPR-FPR is chosen as the fold enrichment cutoff. In mitochondrial and ER-associated APEX-RIP experiments, ROC analysis was based on fold enrichment values; in the nuclear-cytoplasmic partitioning experiment, it was based on calculated nuclear preference scores.

Nuclear preference scores were calculated as follows. To each gene, *i,* we assigned coordinates, (x,y) = (log_2_*N_i_*,log_2_*C_i_*), where *N_i_* and *C_i_* denote to the fold-enrichments from nuclear- (NLS) and cytosolic- (NES) APEX-RIP, respectively. In this space, the line y=x (or rather,log_2_*N* =log_2_*C*) corresponds to all genes that were equally enriched in both nuclear and cytosolic APEX-RIP, and hence do not preferentially reside in either compartment. The nuclear preference score for gene *i* (NPS*i*), is therefore defined as the minimum distance between its coordinates and the line log_2_*N* =log_2_*C*. This is equivalent to calculating the distance between points (x_1_,y_1_) = (log_2_*N_i_*, log_2_*C_i_*) and (x_2_,y_2_) = (0.5(log_2_*N_i_*+log_2_*C_i_*), 0.5(log_2_*N_i_*+log_2_*C_i_*)). Hence:

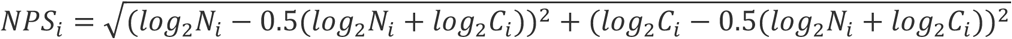

…which reduces to:

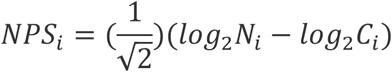

The true and false positive gene sets needed for ROC analysis were defined as follows:

1. For mitochondrial APEX-RIP, true positives corresponded to the thirteen mitochondrial-encoded mRNAs; false positive RNAs corresponded to nuclear-encoded long non-coding RNAs.
2. For the nuclear and cytosolic partitioning experiment, true and false positive gene lists were compiled using available ENCODE human cell line (NHEK-Normal Human Epidermal Keratinocytes) nuclear–cytoplasmic fractionation data(Dunham et al. 2012). We calculated fold-enrichments for RNAs in each compartment (scaled relative to the whole cell RNA, Figure S2A), and used these values to derive Nuclear Preference Scores, as described above. True positive and true negative nuclear RNAs were then defined as the 1000 transcripts with the highest and lowest NPSs, respectively (Figure 2B; Table S2 tab 4). Using these gene lists to perform ROC analysis on the original ENCODE data produced a significance threshold cutoff at an NPS of 1.107 (Figure S2B–D), and lists of the 5467 and 10130 RNAs called as being enriched in the nucleus and cytoplasm, respectively (Table S2, tabs 4 and 5).
3. For ER-APEX-RIP, most true positive genes were defined using data from ER-localized proximity-dependent ribosome profiling(Jan, Williams, and Weissman 2014), corresponding to all RNAs with input RPKM ≥ 5.0, input count ≥ 12, and log2(fold enrichment) ≥ 0.904 (Table S3 tab 4). Additional true positive genes were predicted by Phobius as having secretory signals, and were absent in MitoCarta(Pagliarini et al. 2008). (Rhee et al. 2013)False positive RNAs included all genes lacking secretory signals, as predicted by Phobius, SignalP, and TMHMM.

### Coverage and Specificity analysis of nuclear, cytosolic, and ER-proximal RNAs

To estimate the coverage (recall) and specificity of APEX-RIP at each subcellular compartment, we assembled lists of established target and off-target genes tailored for that compartment.

For analysis of the nuclear–cytosolic datasets (Figure 2F), our reference nuclear gene list comprised 827 lncRNAs with average RNA pre-enrichment abundances of 1.0 or greater. Our reference off-target list comprised the set of 1260 ER-proximal RNAs defined using proximity-restricted ribosome profiling(Jan, Williams, and Weissman 2014).

For analysis of the ER-proximal dataset (Figure 3I–J), our reference gene list comprised 71 mRNAs encoding ER-resident proteins (Table S3 tab 5). Our reference off-target list comprised (7589) RNAs lacking secretory annotation, as assessed using Phobius(Käll, Krogh, and Sonnhammer 2004), TMHMM(Krogh et al. 2001), SignalP(Petersen et al. 2011), and which lacked the GOCC terms “Endoplasmic reticulum,” “Golgi,” “membrane,” and “extracellular”(Ashburner et al. 2000).

For analysis of contaminants in ER datasets (Figure S3E), the RNAs that encode proteins with no predicted secretory annotation by Phobius, TMHMM, and SignalP and lacked GOCC terms “Endoplasmic reticulum,” “Golgi,” “membrane,” and “extracellular” were submitted to DAVID Bioinformatics analysis(Huang, Sherman, and Lempicki 2009) to find Gene ontology term enrichment against human background. Only terms with p-values less than 0.05 were shown.

---

**Supporting Figure 1.**
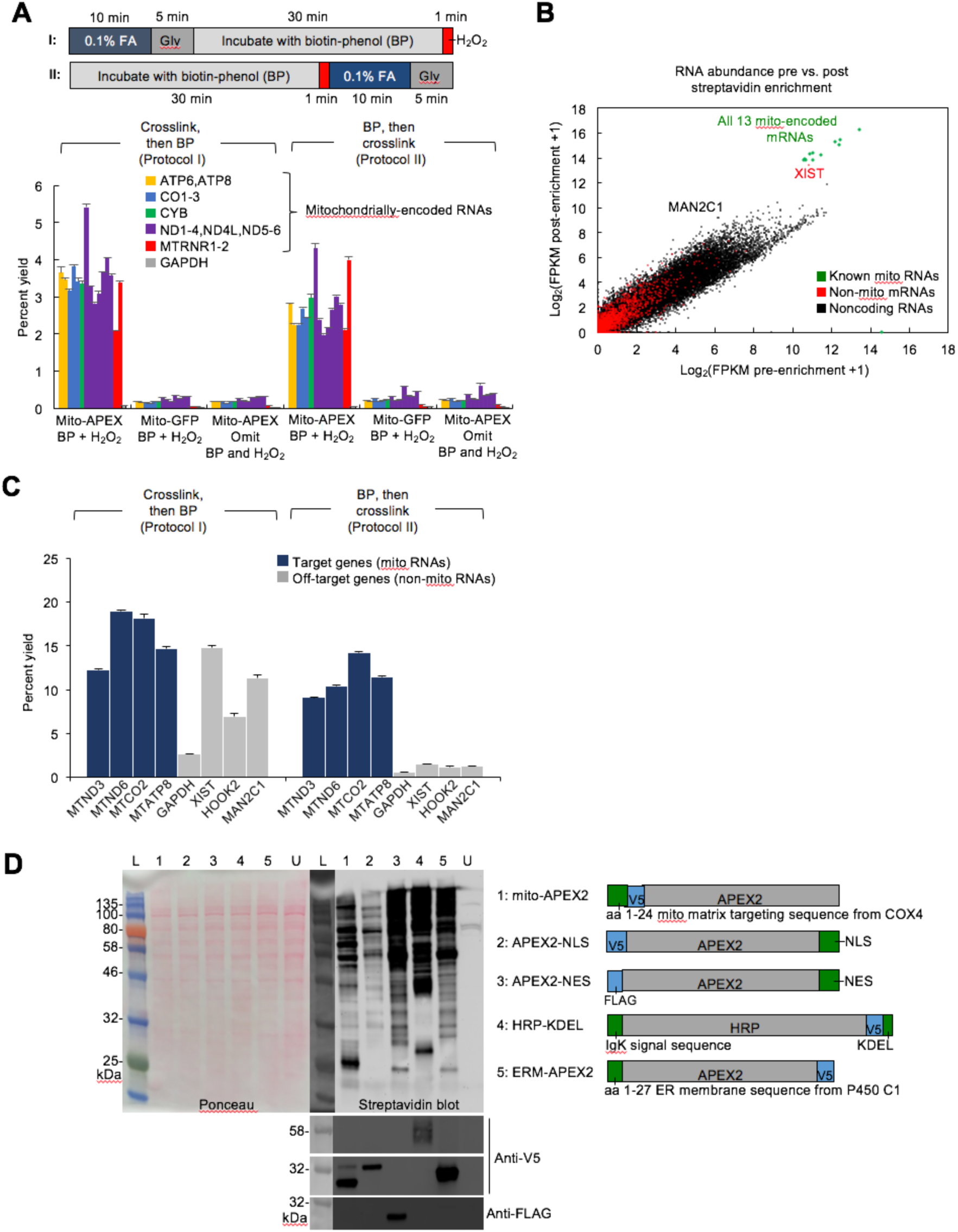
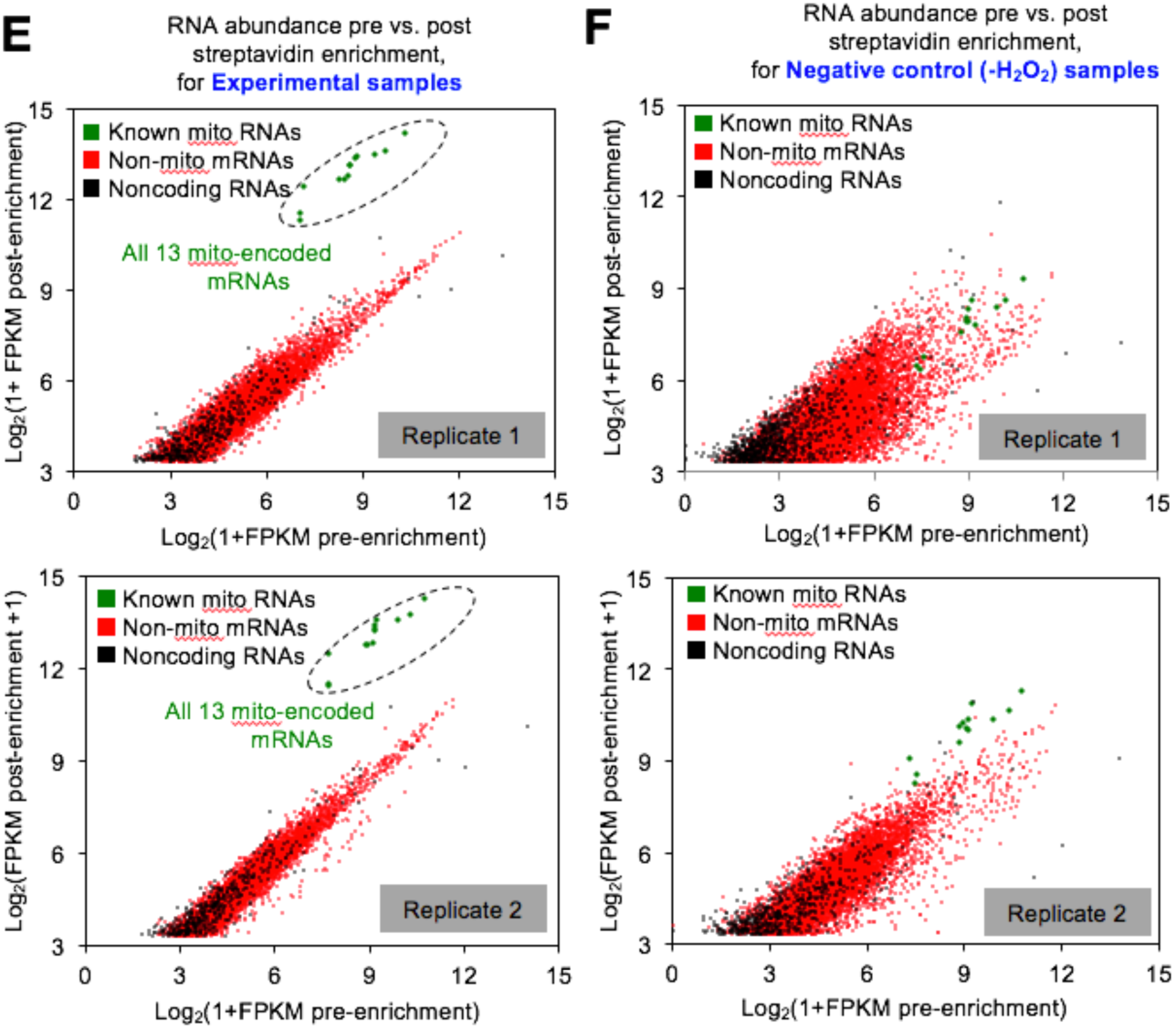
Optimization of APEX-RIP protocol and additional mitochondrial APEX-RIP data (related to **Figure 1**). (**A**) *Top:* Alternate labeling and crosslinking protocols. In protocol I., cells are crosslinked with formaldehyde (FA) and quenched with Glycine (Gly) prior to the introduction of biotin-phenol (BP) and the initiation of APEX-catalyzed biotinylation with H_2_O_2_. In Protocol II., live cells are incubated in BP, and *in situ* biotinylation is initiated prior to FA crosslinking. In both cases, cell lysis, streptavidin enrichment and RNA purification proceed as described (*see methods*). *Bottom:* qRT-PCR analysis comparing Protocols I and II. Negative control experiments replace mito-APEX with mito-GFP, omit BP or omit H_2_O_2_. All constructs were APEX1 derivatives, transiently expressed in HEK 293T cells. Data are the means of three replicates ± one standard deviation. (**B**) RNA-Seq analysis of RNAs enriched by protocol I. Although all 13 mitochondrially-encoded mRNAs (green) were enriched, these were accompanied by substantial contaminating RNAs, including *XIST* and *MAN2C1*. (**C**) qRT-PCR analysis comparing Protocols I. and II., including off-target controls designed using the results from RNA-Seq. Note the superior enrichment obtained using Protocol II. Cells in this experiment stably-expressed mito-APEX2. This protocol and cell line were used to collect all data in (Figure 1). Data are the means of three replicates ± one standard deviation. (**D**) Characterization of all APEX2 fusion constructs used in this work. HEK 293T cells stably expressing the indicated constructs (*right*) were labeled and crosslinked via Protocol II., lysed and analyzed by SDS-PAGE, blotting with streptavidin-HRP, Anti-V5 and anti-FLAG. L: ladder; U: untransfected HEK 293T cells. (**E**) Data from additional replicates of the mitochondrial APEX-RIP experiment, depicted as in (Figure 1D). (**F**) In the absence of H_2_O_2_ treatment, mito-APEX-RIP fails to enrich mitochondrial target genes. Data from individual replicates are shown.

**Supporting Figure 2.**
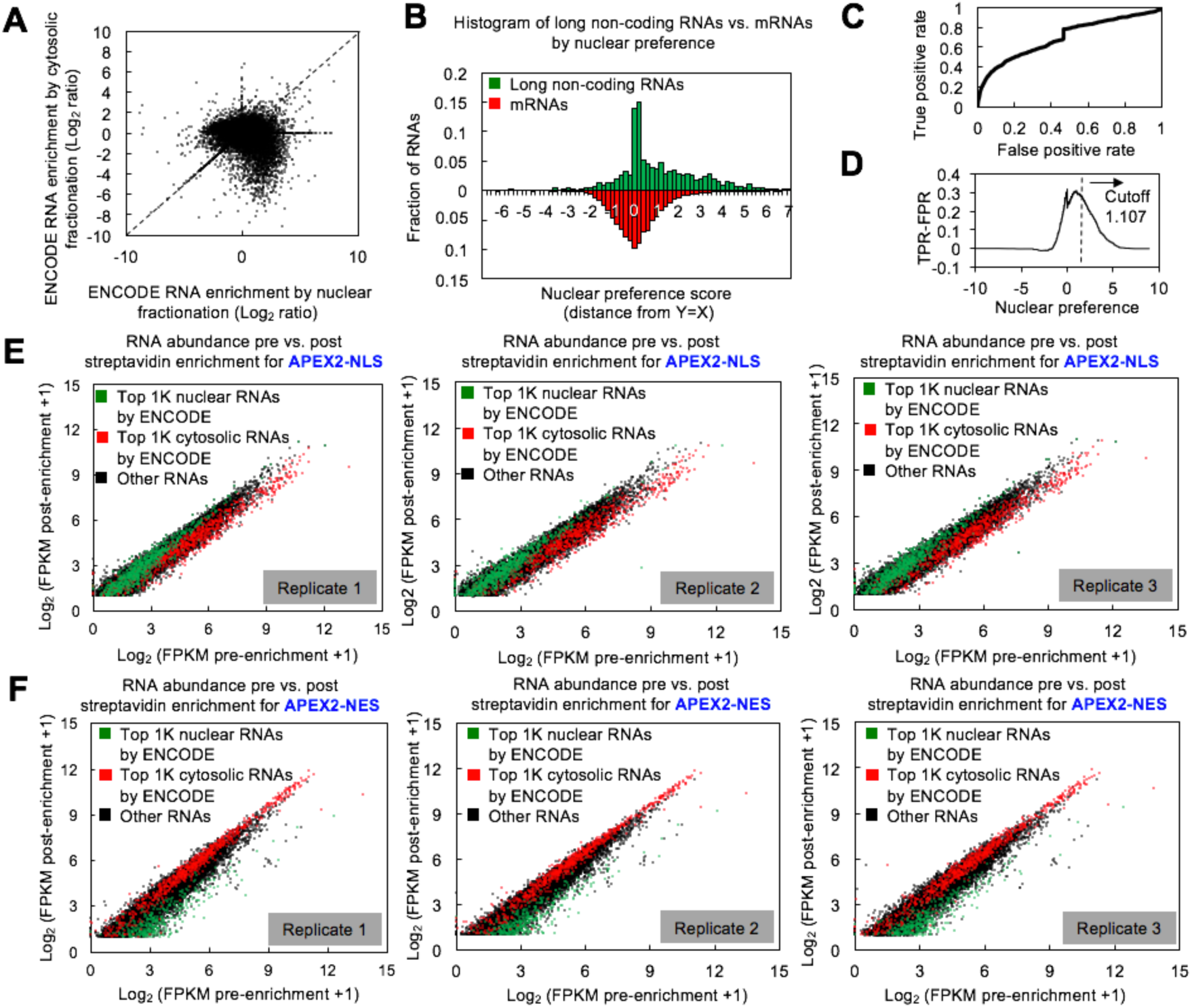
Identification of ENCODE-derived nuclear RNA standards; additional representations of the APEX-RIP nuclear—cytoplasmic data. (**A**) human cell line (NHEK) fractionation data from ENCODE(Dunham et al. 2012), illustrating the relative enrichment of genes during nuclear and cytosolic fractionation. RNA enrichments are calculated as in (Figure 2B) (*see methods*). Displayed are the average values of three replicates, for all genes with FPKM_pre enrichment_ ≥ 1.0. Dashed line denotes y = x line. (**B**) Distribution of genes by nuclear preference score (*see methods*). lncRNAs, predicted to be mostly nuclear, are plotted in green, and mRNAs, predicted to be mostly cytosolic, are plotted in red. (**C–D**) Selection of nuclear-enriched RNA standards using ENCODE fractionation data. (**C**) ROC analysis applied to histogram in (B). For each nuclear preference cutoff value, the True Positive Rate (TPR)–defined as the fraction of lncRNAs above the cutoff–was plotted against the False Positive Rate (FPR)–defined as the fraction of mRNAs above the cutoff. (**D**). Using the output of ROC analysis, a nuclear preference score cut-off value was calculated, defined as the first local maximum in the graph of (TPR-FPR) versus Nuclear Preference Score. This value was applied to obtain a list of 3056 nuclear-enriched standard RNAs. (**E**) Data from individual replicates of the NLS-APEX-RIP experiment. Genes enriched by nuclear and cytoplasmic fractionation in the ENCODE dataset are colored green and red, respectively. (**F**) Data from individual replicates of the NES-APEX-RIP experiment, depicted as in (**E**).

**Supporting Figure 3.**
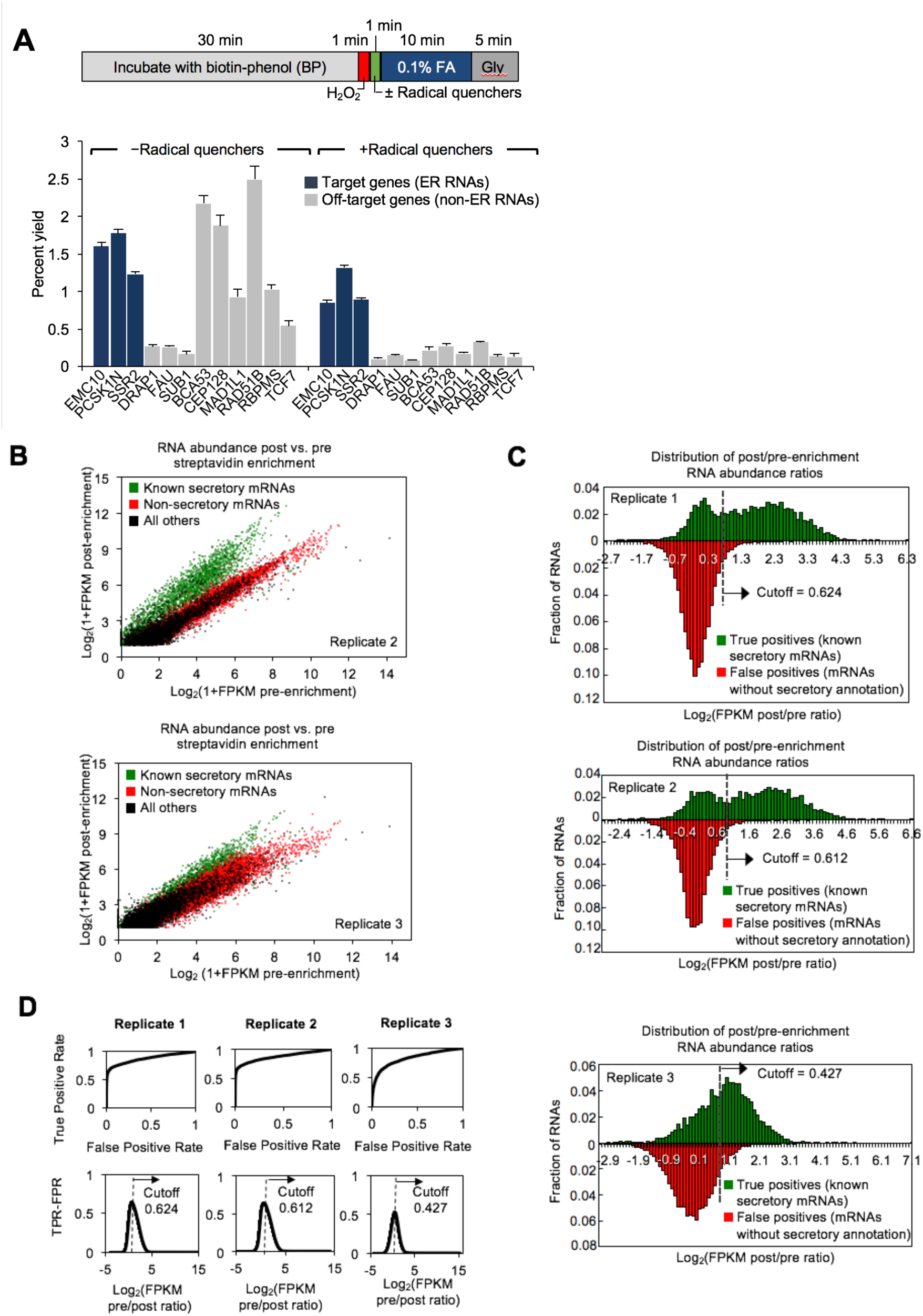
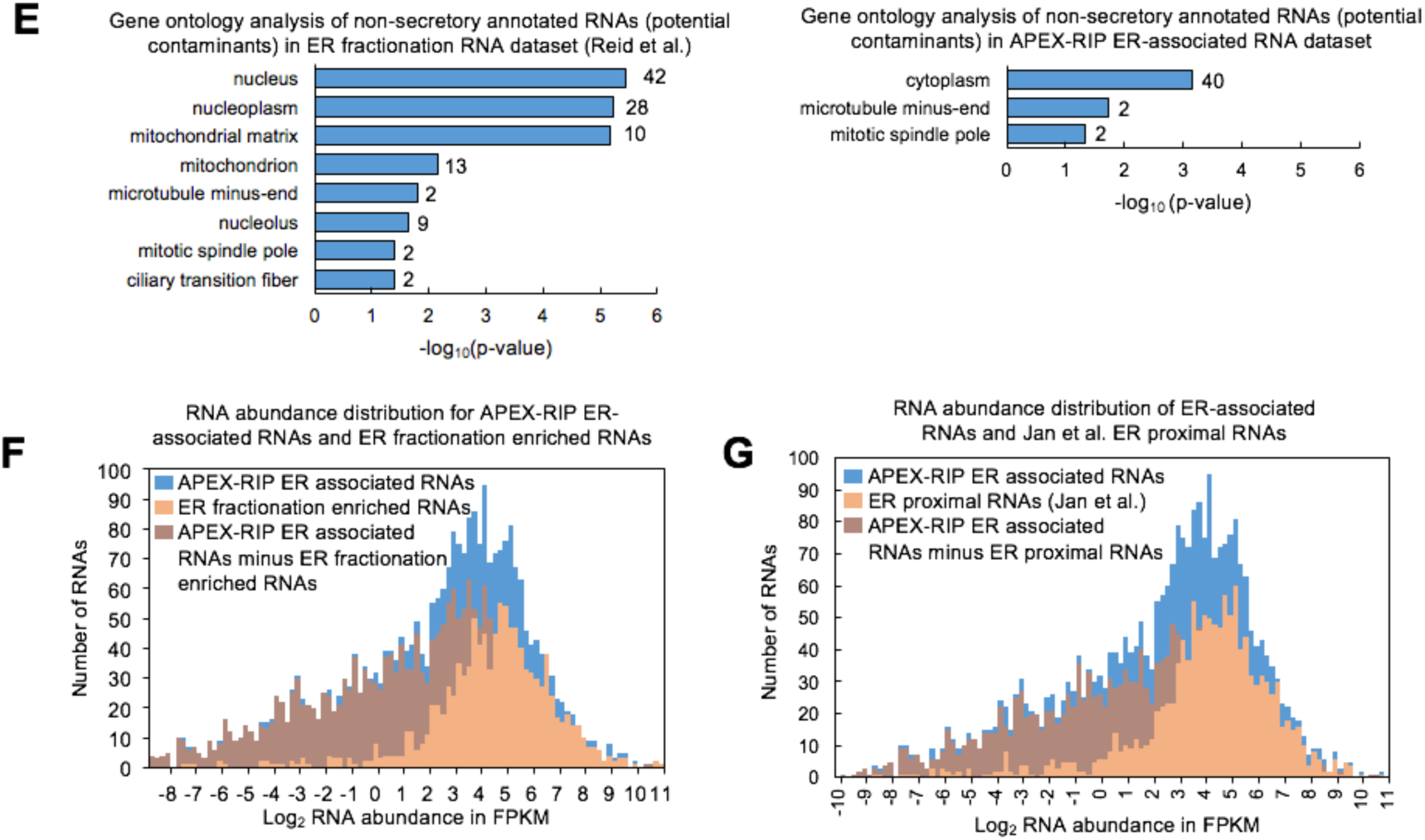
Further optimization of the APEX-RIP protocol; additional HRP-KDEL data (Related to **Figure 3**). (**A**) Addition of a radical quenching step between APEX2 labeling and formaldehyde crosslinking improves the specificity of RNA capture. *Top:* schematic of the revised labeling–crosslinking workflow. *Bottom:* qRT-PCR analysis of the ER-APEX-RIP experiment with and without this additional quenching step, as in (Figure S1C). The radical quenchers used were Trolox and ascorbic acid. (**B–D**) Quality assessment for individual replicates of the ER-APEX-RIP experiment. (**B**) Data for two additional replicates of the HRP-KDEL APEX-RIP experiment, depicted as in (Figure 3F). (**C**) Histograms showing the distribution of RNA enrichment values–log_2_([FKPM _post enrichment_]/[FKPM _pre enrichment_])–for mRNAs encoding known secretory proteins (top histograms, green), and mRNAs encoding non-secretory proteins (bottom histograms, red). Significance cutoffs were determined using ROC analysis (*see below*). Known secretory and non-secretory standard mRNAs were cataloged as described (*see methods*). (**D**) Determination of significance thresholds ratio cut-offs by ROC analysis. Data were processed as described in Figures S2D–E, using True Positive secretory and False Positive non-secretory standard RNAs (*see methods*). Given these analyses, and those in (B–C), only data from Replicates 1 and 2 were used to generate our final ER-associated RNA list. To do this, the individual ROC-derived thresholds were used to calculate significantly enriched genes from each dataset, and the final ER-associated RNA list was defined as the intersection of these two lists. (**E**) Gene ontology (GO) analysis of mRNAs in ER datasets lacking secretory annotation. *Left*: non-secretory mRNAs enriched by biochemical fractionation (Reid and Nicchitta 2012) predominantly exhibit nuclear and mitochondrial annotation. *Right*: non-secretory mRNAs enriched by ER-APEX-RIP have predominantly cytosolic annotation. All terms shown have *p* < 0.05, as assessed using DAVID(Huang, Sherman, and Lempicki 2009). (**F—G**) APEX-RIP recovers low-abundance targets with greater efficiency than do conventional approaches. In each case, starting abundances are defined by ENCODE. (**F**) Comparison of APEX-RIP to biochemical fractionation. (**G**) Comparison of APEX-RIP to proximity-restricted ribosome profiling.

**Supplementary Table S1:**
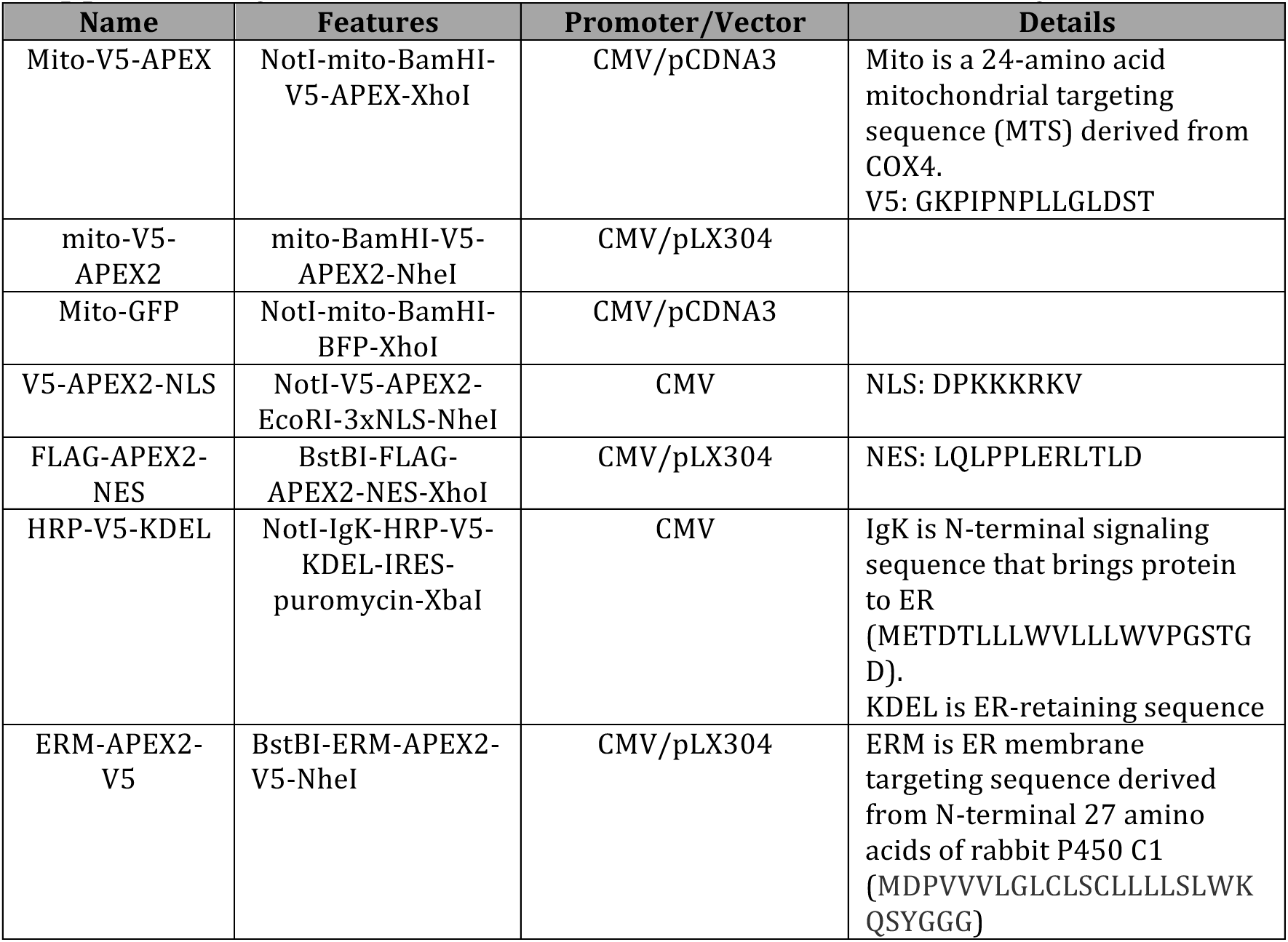
Genetic constructs used in this study.

**Supplementary Table 2.** Antibodies used for immunofluorescence.

| Antibody | Source | Company | Catalog number | Dilution | Associated Figure |
| --- | --- | --- | --- | --- | --- |
| Anti V5 | Mouse | Life Technologies | R960-25 | 1:1000 | Figure 1B, 2A, and 3A |
| Anti FLAG | Mouse | Agilent | 200472 | 1:500 | Figure 2A |
| Anti Mouse-AlexaFluor488 | Goat | Life Technologies | A-11029 | 1:1000 | Figure 1B, 2A, and 3A |

**Supplementary Table 3.**
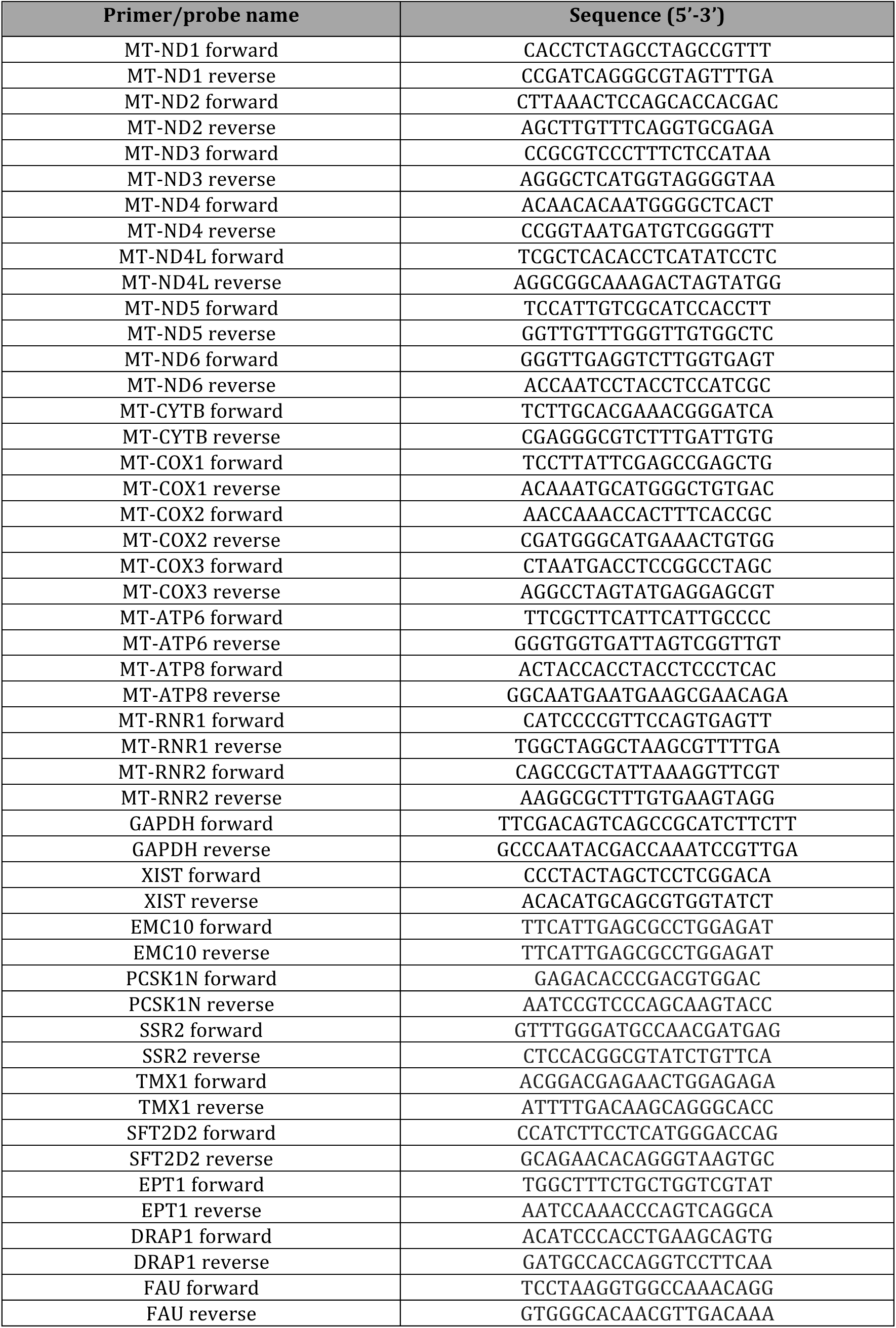
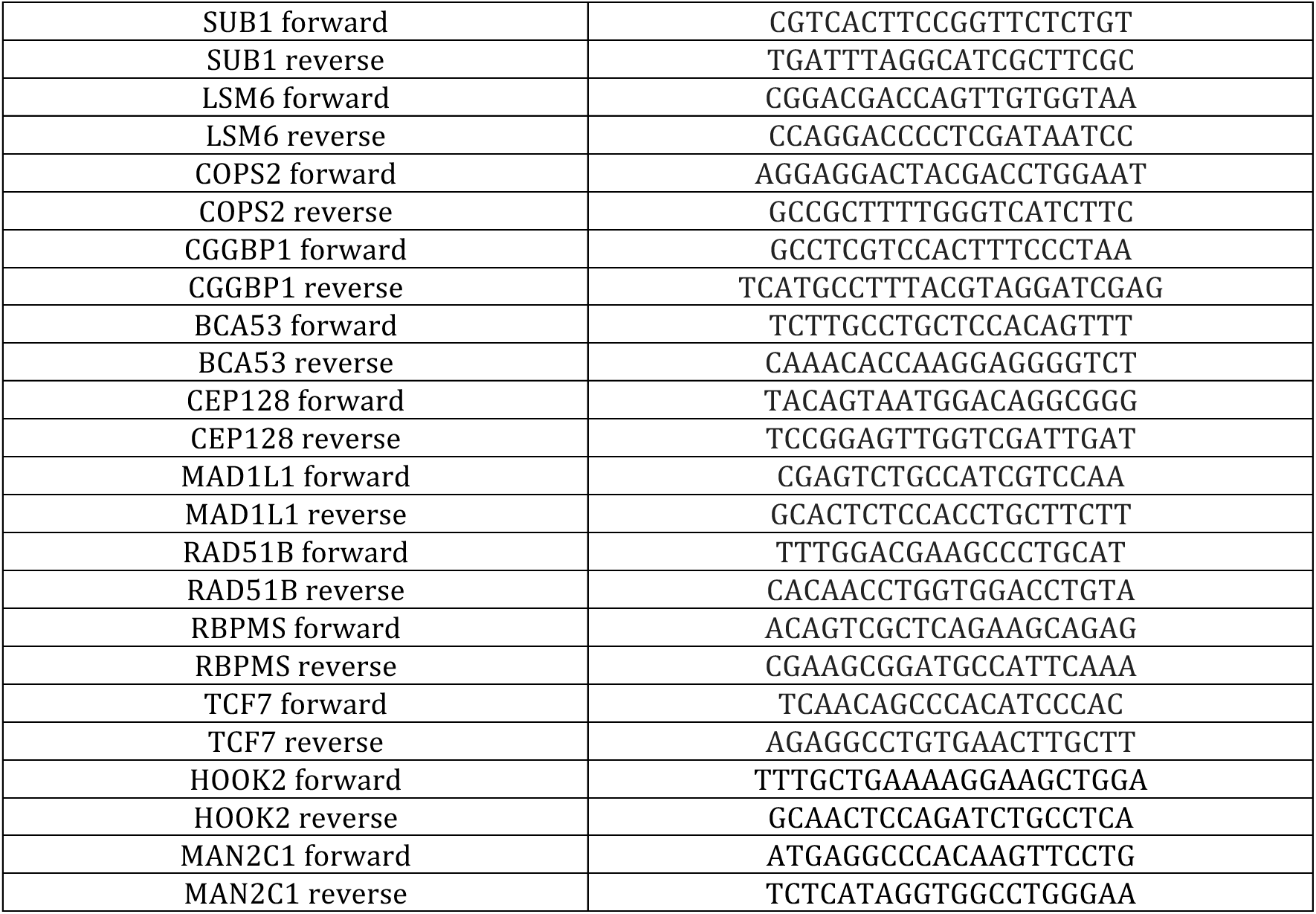
qRT-PCR primers used in this study.

